# Intracellular delivery of full-length antibodies via organ targeted lipid nanoparticles

**DOI:** 10.1101/2025.09.26.678781

**Authors:** Azmain Alamgir, Militsa Yaneva, Mor Sela, Patricia Mora-Raimundo, Souvik Ghosal, Anas Odeh, Peleg Hasson, Rory C. Chien, Avi Schroeder, Matthew P. DeLisa, Christopher A. Alabi

## Abstract

Antibodies are proteins prized for their ability to bind to extracellular antigens with exceptionally high affinities and specificities. These features have motivated researchers to utilize antibody-antigen binding to inhibit intracellular disease targets in the proteome, yet delivery of antibodies into the cytosol of cells has long been a considerable challenge. Here, we outline the development of a novel lipid nanoparticle (LNP) platform for delivering antibodies into cells to selectively inhibit disease-relevant cytosolic targets. This approach efficiently delivers various therapeutic antibodies into multiple cancer cell lines, inhibiting key transcription factors in inflammatory and cancer signaling pathways. We further demonstrate systemic delivery of therapeutic antibodies in disease models, including alpha-synuclein-specific antibodies for Parkinson’s disease and RelA-specific IgGs for acute lung injury using targeted LNP formulations. This work establishes a promising method for using LNPs for the delivery of antibody and antibody-derived therapeutics intracellularly to treat numerous proteome targets.

## Introduction

Immunoglobulin (IgG) antibodies, best known for their pivotal role in the body’s immune system, have emerged as one of the most important therapeutic modalities in the biotechnology industry over the past several decades. Since the introduction of the first antibody therapeutic in 1986,^1^ over 100 FDA-approved monoclonal antibodies (mAbs) have made their way into the clinic for treatment of ailments ranging from rheumatoid arthritis, chronic bowel inflammations, and numerous different cancers.^2^ In 2022 alone, mAbs comprised over half of all drugs gaining FDA approval and represented ∼80% of the total market revenue from protein-based biopharmaceuticals.^3^ Antibody therapies possess high serum stabilities, prolonged circulation lifetimes, and perhaps most importantly, exceptional specificities against target antigens.^4^ Combined with established and scalable discovery pipelines (hybridoma technology^5,6^ and surface display platforms^7,8^), antibodies possess key advantages that make them a compelling modality to treat degenerative diseases as compared to traditional small molecule drugs and emergent nucleic acid-based therapies.^9,10^

Virtually all antibody therapeutics act on extracellular antigens by binding to soluble proteins or cell-surface receptors. The reason antibody development efforts have focused exclusively on extracellular targets is due to the inability of these large, globular macromolecules to translocate across cell membranes. However, due to their high affinity and specificity, there is increasing interest in using antibodies for drugging protein targets that reside within cells.^11–17^ A large motivation for utilizing antibodies with intracellular modes of action is to inhibit the “undruggable” proteome – a collection of proteins that comprise over 80% of the total human proteome which lack well-defined binding sites and thus are intractable using conventional small molecule therapeutics.^18^ Hence, expanding the accessibility of antibodies to intracellular protein antigens could open the door to novel therapeutic applications against undruggable targets while also streamlining their use as basic research tools for dissecting cell pathology and signaling.

Introducing antibodies into cells requires delivery methods that can effectively transport them across cell membranes, which as previously alluded to, poses a significant challenge due to the membrane impermeability of large, globular proteins. While physical membrane disruption methods such as microinjection^19,20^ and electroporation^21^ can be used to deliver antibodies into cells, these methods suffer from low throughput and incompatibility *in vivo*. Carrier-mediated approaches, such as polymeric assemblies,^13^ lipid-based vehicles,^12,15^ and inorganic nanostructures,^22,23^ as well as engineered antibody conjugates^24,25^ have also been explored for delivering antibodies into cells and are better suited for *in vivo* applications. However, to date, none of these strategies have been clinically validated and therefore face uncertain development paths. Thus, a compelling alternative strategy for antibody delivery is to co-opt delivery methods proven successful for other biological drugs.

To this end, we explored the adoption of clinically validated lipid nanoparticle (LNP) formulations to achieve highly efficient antibody delivery into cells and tissues. LNPs represent one of the most advanced non-viral methods for nucleic acid delivery that have been approved for human use, as exemplified by the success of Patisiran,^26^ the first FDA-approved siRNA therapeutic, and the COVID-19 mRNA vaccines.^27^ Here, we leveraged a bioconjugation-based “cloaking” platform that remodels the surface charge of antibody cargos with cleavable anionic linkers that enable electrostatics-driven assembly with cationic lipids to form LNPs.^28^ We utilized this strategy to functionally deliver off-the-shelf antibodies to the cytosol of living cells, where they selectively inhibited key transcription factors implicated in disease-relevant signaling pathways (**Fig. 1a**). We validated the *in vivo* potential of the delivery platform by successfully delivering alpha-synuclein-specific antibodies to the brains of mice with Parkinson’s disease (PD) and further demonstrated its therapeutic application using organ-targeted LNPs to effectively deliver NF-κB-specific antibodies in a mouse model of acute lung injury (ALI). Taken together, our findings demonstrate the ease and versatility of our cloaking platform for repurposing pre-existing antibodies for targeted protein inhibition following LNP-mediated delivery both *in vitro* and *in vivo*, thereby paving the way for future translation of novel antibody therapies that act on intracellular targets.

**Figure 1.**
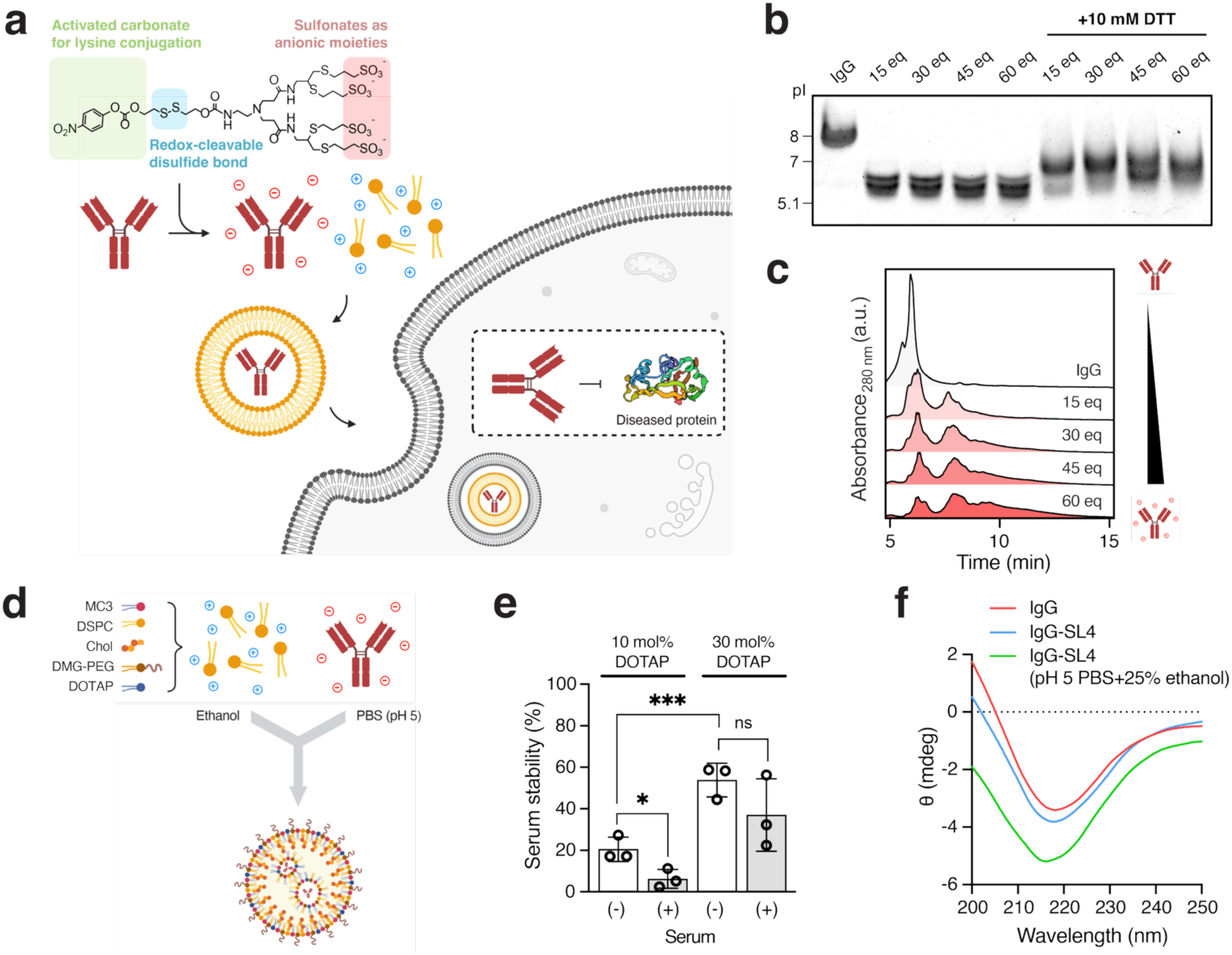
Anionic cloaking of IgG antibodies promotes global surface charge modification and efficient encapsulation into MC3 LNPs. (a) Schematic of anionic cloaking strategy for enabling intracellular IgG delivery using cationic lipids. Chemical modification of surface-exposed lysines with sulfonated cloaking reagents allows electrostatic complexation with cationic lipids to form LNPs. (b) Isoelectric focusing gels of murine isotype IgG samples conjugated to sulfonated compounds, before and after treatment with 10 mM DTT. (c) Chromatograms (measured at 280 nm) from anion exchange chromatography of cloaked IgG samples. (d) Schematic of LNP formulation protocol with cloaked IgGs. Cloaked IgGs are rapidly mixed with lipid components in an aqueous phase consisting of PBS shifted to pH 5. (e) Encapsulation efficiency of cloaked IgG samples (modified with 30 molar eq. of SL4) before and after incubation with mouse serum for 1 hour. MC3 LNPs (MC3/IgG, 2 wt/wt) were formulated with 10 and 30 mol% and 30 mol% DOTAP. Data presented as mean of biological replicates (*n =* 3 for encapsulation efficiency) ± SD. Statistical significance was determined by unpaired t-tests followed by Bonferroni-Dunn correction for multiple comparisons (*p < 0.05, **p < 0.01, ***p < 0.001, ****p < 0.0001). (f) CD spectra of native IgG and cloaked IgG samples. Cloaked IgG samples were placed in pH 5 buffer containing 25% ethanol to mimic formulation conditions (green). Spectra are representative of biological replicates (*n* = 3).

## Results

### Anionic cloaking results in efficient antibody surface charge modification and encapsulation in LNPs

Conventional LNP formation relies on electrostatic interactions between anionic nucleic acid cargos and cationic ionizable lipids in LNP formulation mixtures. To harness the same electrostatics-driven LNP formation with antibody cargos, we utilized an anionic “cloaking” platform that relies on surface charge remodeling of proteins through activated carbonate compounds. These cloaking reagents enable chemoselective attachments of sulfonate groups to surface-exposed lysine residues and contain a disulfide bond that can undergo redox-mediated cleavage and self-immolation to tracelessly regenerate the native antibody within the reducing environment of the cytosol. From our previous studies, we use a tetrasulfonated linker, termed SL4 (**Supplementary Figure S1 and S20-S21**), that has been shown to be effective in enabling LNP-mediated delivery of proteins such as sfGFP, RNase A, and monoclonal antibodies.^28^

We first verified anionic surface charge modification using SL4 on a murine isotype IgG antibody. Efficient charge remodeling was verified by isoelectric focusing, which showed a clear reduction in protein isoelectric point (pI) when the antibody was reacted with increasing molar equivalents of SL4 (**Fig. 1b**). The pI of the antibody reached a plateau when it was reacted with more than 15 molar equivalents of SL4, likely because residual surface exposed lysine residues are rendered inaccessible due to repulsion from the negatively charged surface. Incubation of SL4-modified IgG samples with 10 mM dithiothreitol (DTT), corresponding to the approximate concentration of reducing agent in the cytoplasm,^29^ resulted in an increase in protein pI towards that of the native IgG, indicating successful linker cleavage and immolation. Anionic surface charge modification was further resolved through anion exchange chromatography, which clearly demonstrated a shift in antibody samples modified with increasing amounts of SL4 towards later retention times (**Fig. 1c**). Interestingly, analysis of both native and cloaked IgG samples incubated with 10 mM DTT showed similar retention times in anion exchange chromatograms, suggesting the formation of reduced antibody fragments following disulfide bond reduction (**Supplementary Figure S2**). Ultimately, we determined that reacting with 30 molar eq. of SL4 was sufficient for IgG surface charge remodeling, which was corroborated by previous experiments. ^28^

Next, we investigated the LNP encapsulation efficiency of cloaked IgGs. Although LNPs with nucleic acid cargos are typically formulated with four lipid components, we previously found that supplementation of the conventional LNP formulation with additional permanently cationic lipids such as 1,2-dioleoyl-3-trimethylammonium-propane (DOTAP) was necessary for efficient LNP encapsulation with cloaked protein cargos.^28^ Therefore, LNPs used in this study were formulated using DOTAP together with the “gold standard” ionizable lipid DLin-MC3-DMA (MC3), which is the principal lipid component found in the FDA-approved siRNA drug Patisiran.^26^ Along with MC3, the formulation also included distearoylphosphatidylcholine (DSPC), cholesterol, and 1,2-dimyristoyl-rac-glycero-3-methoxypolyethylene glycol-2000 (DMG-PEG-2000) at a molar ratio of 50/10/38.5/4.5, with samples using either 11.1 or 45 mol/mol (i.e. 10 or 30 mol%) supplemental DOTAP as the fifth component. For protein encapsulation, the cloaked IgG samples were formulated in an aqueous phase at pH 5 by rapidly mixing with the five lipid components that were dissolved in ethanol, keeping the MC3/IgG wt/wt ratio fixed at 2:1 in all samples (**Fig. 1d**).

We investigated MC3 LNP formulations consisting of 10 mol% DOTAP and 30 mol% DOTAP and assessed these formulations for encapsulation efficiency and stability in mouse serum. We observed an increase in IgG encapsulation using MC3 LNPs formulated with a higher mol% DOTAP (consistent with our previous reports^28^), with 30 mol% DOTAP formulations showing over 60% IgG encapsulation and retaining nearly 40% encapsulated IgGs after incubation in serum (**Fig. 1e** and **Supplementary Figure S3**). Thus, we opted to utilize MC3 LNPs supplemented with 30 mol% DOTAP for subsequent IgG delivery experiments. Analysis of 30 mol% DOTAP formulations by transmission electron microscopy (TEM) verified uniform particle morphology with diameters of ∼200 nm (**Supplementary Figure S4a**). Further TEM imaging of particles formulated with cloaked, Nanogold®-labeled IgGs revealed successful encapsulation of IgGs within the interior of LNPs, with an average of one labeled antibody encapsulated per particle (**Supplementary Figure S4b**).

A requisite step for functional antibody delivery is the ability of the antibody cargo to retain its structure following cloaking and formulation into LNPs. To investigate these effects, we analyzed the secondary structure of IgGs before and after cloaking by circular dichroism (CD) analysis. Overall, we found that cloaking caused only minimal perturbations to antibody structure as evidenced by negligible shifts in measured ellipticity (**Fig. 1f**). We also analyzed structure of cloaked IgG samples placed in pH 5 solution containing 25% ethanol, which was intended to mimic the formulation conditions under which LNPs associate with cloaked proteins (**Fig. 1f** and **Supplementary Figure S5**). Reassuringly, CD measurements confirmed that IgG structure remained mostly undisturbed under these conditions, with only a slight decrease in ellipticity that was indicative of β-sheet formation due to possible aggregation.^30^ Taken together, we confirm that cloaking resulted in efficient anionic modification of IgGs with minimal perturbation to secondary structure, thus enabling IgG encapsulation in LNPs supplemented with DOTAP with sufficient stability.

### Delivery of cloaked IgGs with cationic lipids is robust in multiple cancer cell lines

We next assessed uptake of LNP-encapsulated IgGs into multiple different tumor cell lines, namely A549 non-small cell lung adenocarcinoma cells, DLD-1 colorectal adenocarcinoma cells, and HepG2 hepatocellular carcinoma cells. For these experiments, 200 nM of cloaked murine isotype IgGs was formulated in either MC3 LNPs as described above (**Supplementary Table 1**) or Lipofectamine 2000 (LF2K), a commercial cationic lipid transfection reagent. The uptake of fluorescently-labeled cloaked IgG samples into the three cell lines was highly efficient as quantified by flow cytometric analysis. IgGs exhibited transfection efficiencies ranging from 20-60% with LF2K, while transfection efficiencies with MC3 LNPs exceeded 90% (**Fig. 2a**). In contrast, IgGs alone did not internalize into any cells. We also performed transfections of cloaked IgGs encapsulated in LNPs at concentrations ranging from 10 to 500 nM across the tested cell lines. We observed transfection efficiencies exceeding 80% for all cells at concentrations as low as 100 nM (**Fig. 2b**). Notably, the measured mean fluorescence intensity (MFI) of cells continued to increase with increasing concentration of IgGs (**Fig. 2c**), indicating that the total uptake of protein into cells continued to rise even though transfection efficiency plateaued near 100% at concentrations above 200 nM of IgG. LNP-mediated delivery of IgGs was also found to be largely nontoxic, although DLD-1 cells did exhibit close to 40% reduction in viability following treatments with formulations containing the highest concentration of cloaked IgG (**Fig. 2d**).

**Figure 2.**
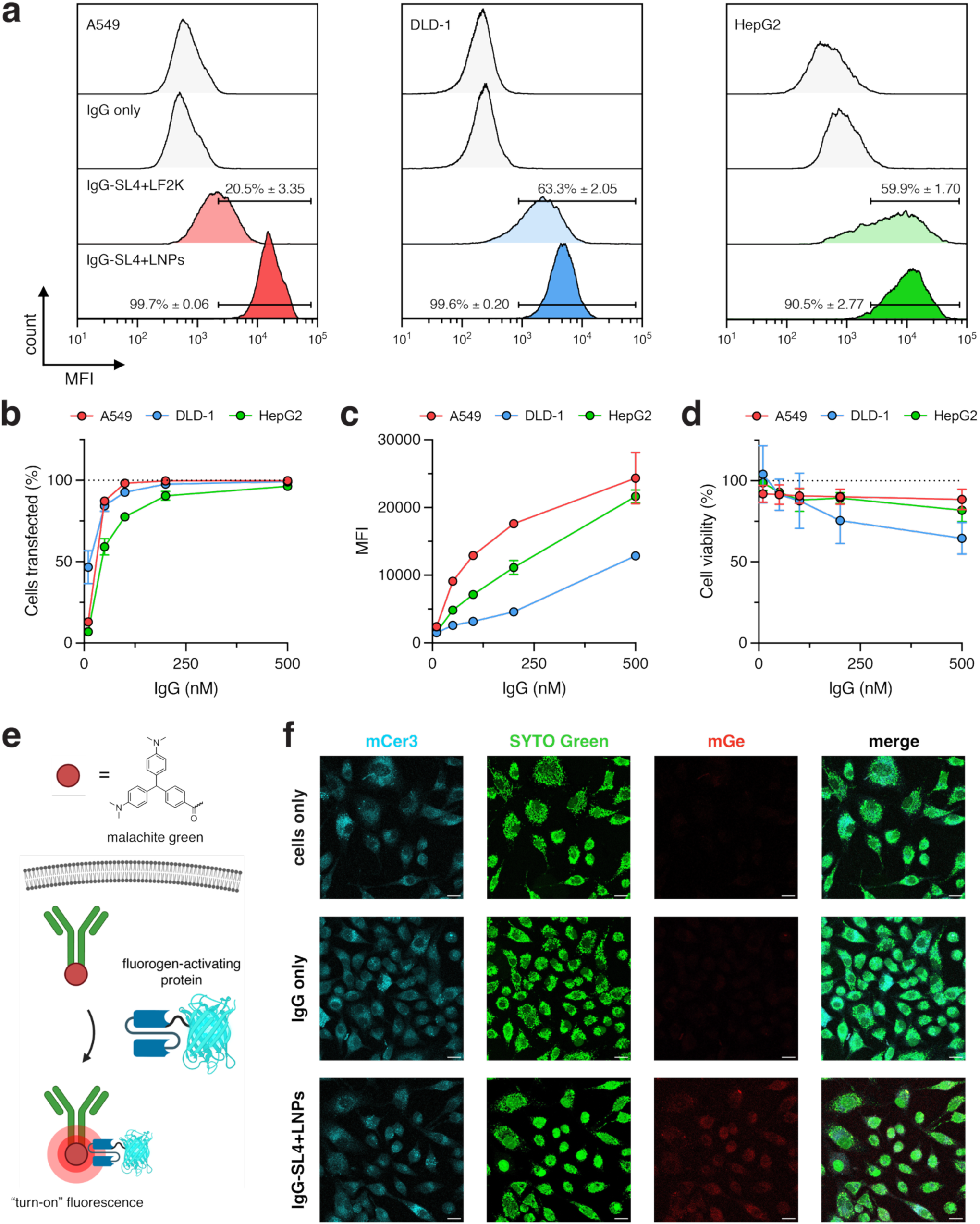
Cellular internalization of cloaked IgGs encapsulated in MC3 LNPs is highly efficient in multiple cancer cell lines. (a) Representative flow cytometry histograms of A549, DLD-1, and HepG2 cancer cells transfected with 200 nM IgGs cloaked with SL4 (30 molar eq.) and encapsulated in MC3 LNPs (MC3/IgG, 2 wt/wt) supplemented with 30 mol % DOTAP. For comparison, cells were transfected with 200 nM of cloaked IgGs complexed with LF2K or IgGs alone. Transfections were performed for 6 hours. Histogram gating to determine percent AlexaFluor 488 labeled IgG-positive cells is overlaid on plots. (b) Percent AlexaFluor 488 labeled IgG-positive cells, (c) mean fluorescence intensity (MFI), and (d) cell viability following transfections performed as in (a) with cloaked IgGs encapsulated in MC3 LNPs. Cell fluorescence was measured by flow cytometry and viability of cells following transfections was measured by MTS assay. Transfections for flow cytometry and cell viability experiments were performed for 6 and 24 hours, respectively. (e) Schematic depicting the fluorogen activating protein (FAP) assay to detect cytosolic delivery of IgGs. (f) Representative confocal microscopy images of A549 cells transfected cloaked IgGs encapsulated in MC3 LNPs or IgGs alone. IgGs were labeled with malachite green isothiocyanate (mGe), and cytosolic IgGs were detected using the FAP assay. Scale bar = 10 μm. All data are the mean of biological replicates (*n* = 3) ± SD.

To achieve functional inhibition, antibodies must be delivered in the cytosol to engage their respective intracellular targets. This is of particular importance given reports estimating the endocytic escape of cargos encapsulated in LNPs to be less than 5%.^31^ To rigorously assess explicit cytosolic delivery of IgG cargo, we utilized a fluorogen activating protein (FAP) assay to quantify how much of the IgG payload was able to escape endosomes following LNP-mediated internalization. For the fluorogen activating protein, we used an anti-malachite green (mGe) single-chain antibody fragment (scFv) fused to mCerulean3 that was expressed in the cytosol. Upon binding to mGe-conjugated IgGs in the cytosol, the FAP constrains the fluorogen and produces a “turn-on” fluorescence signal that is indicative of the amount of LNP-delivered IgG that has successfully reached the cytosol (**Fig. 2e** and **Supplementary Figure S6**).^32^ When A549 cells were co-transfected with the FAP reporter along with LNP-encapsulated cloaked IgGs labeled with mGe, diffuse cytosolic signals were visible by confocal microscopy (**Fig. 2f**). In agreement with the flow cytometry results above, LNPs formulated using 30 mol% DOTAP resulted in stronger FAP-based fluorescence than those formulated with 10 mol% DOTAP (**Supplementary Figure S7**), indicating more efficient cytosolic IgG delivery and providing further evidence that higher DOTAP concentrations not only increase particle stability but also promote more efficient IgG uptake with LNPs.

### Cytosolic delivery of transcription factor-specific IgGs results in effective signaling inhibition

Next, we performed functional inhibition assays to test the inhibitory effects of several commercially available antibodies that bind transcription factors implicated in disease-associated signaling pathways (**Fig. 3a**). We specifically investigated inhibition of several key transcription factors that regulate signaling in the following pathways: (i) NF-κB pathway in A549 cells using murine antibodies that bind to the RelA and p50 components of the heterodimeric NF-κB complex; (ii) Wnt pathway in DLD-1 cells using antibodies that bind to the N- and C-terminus of β-catenin; and (iii) JAK-STAT pathway in HepG2 cells using antibodies that bind STAT3 and its Tyr705 phosphorylated isoform. These pathways were chosen due to their pivotal roles in inflammation and cancers.^33–35^

**Figure 3.**
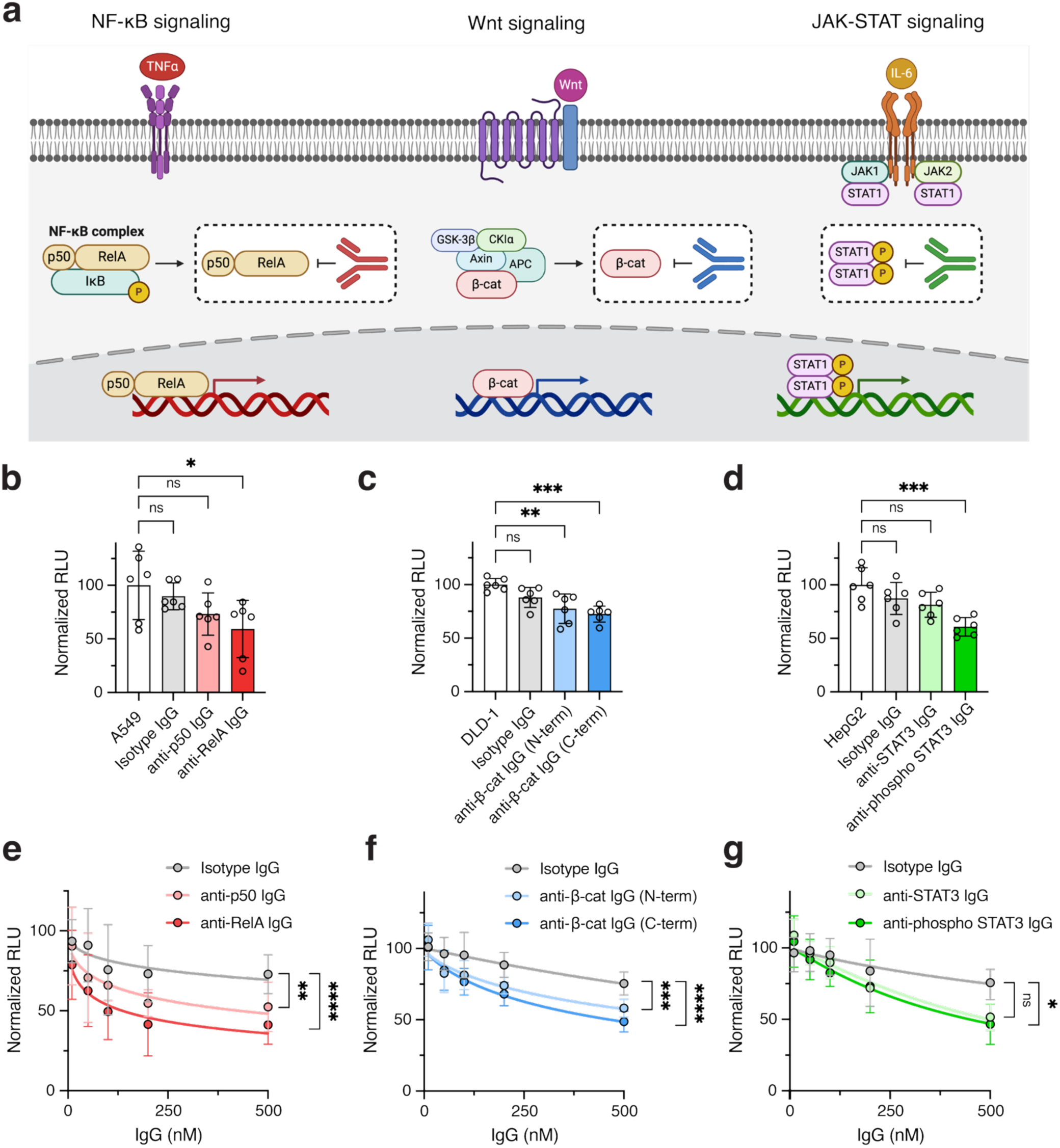
Intracellular delivery of IgGs with cationic lipids results in potent inhibition of multiple signaling pathways *in vitro*. (a) Schematic of investigated cell signaling pathways and transcription factor targets recognized by delivered IgGs. (b-d) Luminescence readouts of transcriptional activity following transfections of 200 nM IgGs cloaked with 30 molar eq. of SL4 and complexed with LF2K. Tested antibodies included: (b) anti-RelA and anti-p50 IgGs against NF-κB signaling in A549 cells, (c) N-terminal and C-terminal anti-β-catenin IgGs against Wnt signaling in DLD-1 cells, and (d) anti-STAT3 and anti-phospho STAT3 IgGs against JAK-STAT signaling in HepG2 cells, along with isotype control IgGs. (e-g) Luminescence readouts of transcriptional activity following transfections of the same cloaked IgGs in (b-d) but encapsulated with MC3 LNPs (MC3/IgG, 2 wt/wt and supplemented with 30 mol % DOTAP; formulated in PBS pH 5). All transfections were performed for 12 hours. All data are mean of biological replicates (*n* = 6) ± SD. Statistical significance was determined by ordinary one-way ANOVA followed by Bonferroni correction for multiple comparisons and two-way ANOVA followed by Bonferroni correction for multiple comparisons (\**p* < 0.05, \*\**p* < 0.01, \*\*\**p* < 0.001, \*\*\*\**p* < 0.0001).

To first verify binding activity of IgGs following anionic cloaking with SL4, we subjected the cloaked anti-RelA antibody to ELISA analysis using purified RelA protein as the immobilized antigen. As expected, binding activity for the anti-RelA IgG was diminished following cloaking but could be effectively restored following additional DTT treatment to remove the cloaking agent (**Supplementary Figure S8**). We next evaluated intracellular binding and signaling inhibition by transfecting the three different cancer cell lines with cloaked IgGs complexed with LF2K. Following transfection, the effect on signaling was evaluated using luciferase-based transcriptional assays. Importantly, each of the tested IgGs exhibited clear target protein inhibition relative to cells treated with a LF2K-complexed isotype IgG control (**Fig. 3b-d** and **Supplementary Figure S9**). The best performing IgGs following LF2K-mediated transfections were the anti-RelA and anti phospho-STAT3 antibodies that each showed nearly 50% reduction in NF-κB signaling (**Fig. 3b**) and STAT3 signaling (**Fig. 3d**), respectively.

Encouraged by these results, we proceeded to generate MC3 LNP formulations of these cloaked antibodies. Following transfection into the same three cancer cell lines, we observed that nearly all the MC3 LNP-encapsulated IgGs triggered statistically significant, dose-dependent decreases in cell signaling across all three pathways compared to MC3 LNP-encapsulated isotype IgG controls (**Fig. 3e-g** and **Supplementary Figure S9**). At the highest IgG doses (500 nM), ∼50-60% inhibition of signaling activity was achieved with each IgG, with the anti-RelA antibody again achieving the most potent inhibition among the tested IgGs. To further confirm target engagement in the cytosol, we performed co-immunoprecipitation experiments. Specifically, A549 cells transfected with MC3 LNP-encapsulated anti-RelA IgGs were lysed, and the resulting lysates were subjected to immunoprecipitation (IP) using Protein G-coated beads to pull down IgG-RelA complexes. Western blot analysis of IP samples using anti-RelA antibody demonstrated successful pull-down of intracellular RelA (**Supplementary Figure S10**), thereby confirming target-specific engagement of a cytosolic target by LNP-delivered antibodies *in vitro*.

Recently, it has been shown that IgGs delivered intracellularly following cell permeabilization can interact with TRIM21, an endogenous E3 ligase that binds to the Fc region of IgG antibodies.^36,37^ Recruitment of TRIM21 to the IgG-bound antigen results in ubiquitination of the antigen and its subsequent degradation by the 26S proteasome. This method, called “Trim-away”, has been leveraged for targeted protein degradation.^36,37^ To explore if the TRIM21 pathway is implicated in antibody-mediated inhibition in our system, we performed TRIM21 knockout experiments with siRNA and investigated the effects on NF-κB inhibition with anti-RelA IgGs. Interestingly, we observed higher reduction of NF-κB transcriptional activity in TRIM21-silenced A549 cells, with more pronounced inhibition occurring at higher amounts of TNF-α stimulation (**Supplementary Figure S11**). These findings suggest that the TRIM21 pathway may be interfering with antibody-induced inhibition via proteosome-dependent antibody degradation.

Overall, the results from these experiments show that pre-existing antibodies, including those that are commercially available, can be repurposed for intracellular inhibition of a diverse range of transcription factors. Further, the data validates that functional inhibition can be achieved using both cationic lipid transfection reagents as well as clinical LNP formulations, thereby paving the path for selective inhibition of intracellular targets using off-the-shelf antibodies for both *in vitro* and *in vivo* experimentation.

### LNP-encapsulated IgGs distribute to major organs *in vivo* following systemic delivery

To assess the *in vivo* delivery capability and biocompatibility of our system, we analyzed biodistribution of systemically injected LNP-encapsulated IgGs. To this end, cloaked IgGs were first labeled with Cy5, encapsulated in MC3 LNPs, and systemically administered into BALB/c mice via tail vein injection (1 mg/kg of total protein). *Ex vivo* fluorescence images of harvested organs revealed minimal accumulation of free IgG primarily to the liver and kidneys, whereas IgGs encapsulated in LNPs showed enhanced uptake in the spleen, liver, and lungs of mice (**Fig. 4a** and **Supplementary Figure S12**). Delivery of MC3 LNP-encapsulated IgGs to the lungs was particularly remarkable, with approximately half of the IgG dose localizing to this organ within 1 hour (**Fig. 4b** and **c**). Extrahepatic delivery of LNPs to the lungs is well known to be modulated by supplementation of traditional LNPs with cationic lipids such as DOTAP,^38,39^ which we also observed with LNP-encapsulated protein cargos.^28^ It is also worth noting that whereas free IgGs become rapidly cleared from circulation after 6 hours post-injection, MC3 LNP-encapsulated IgGs persist in the liver and to a lesser extent in the lung for at least 6 hours (**Fig. 4b** and **c**).

**Figure 4.**
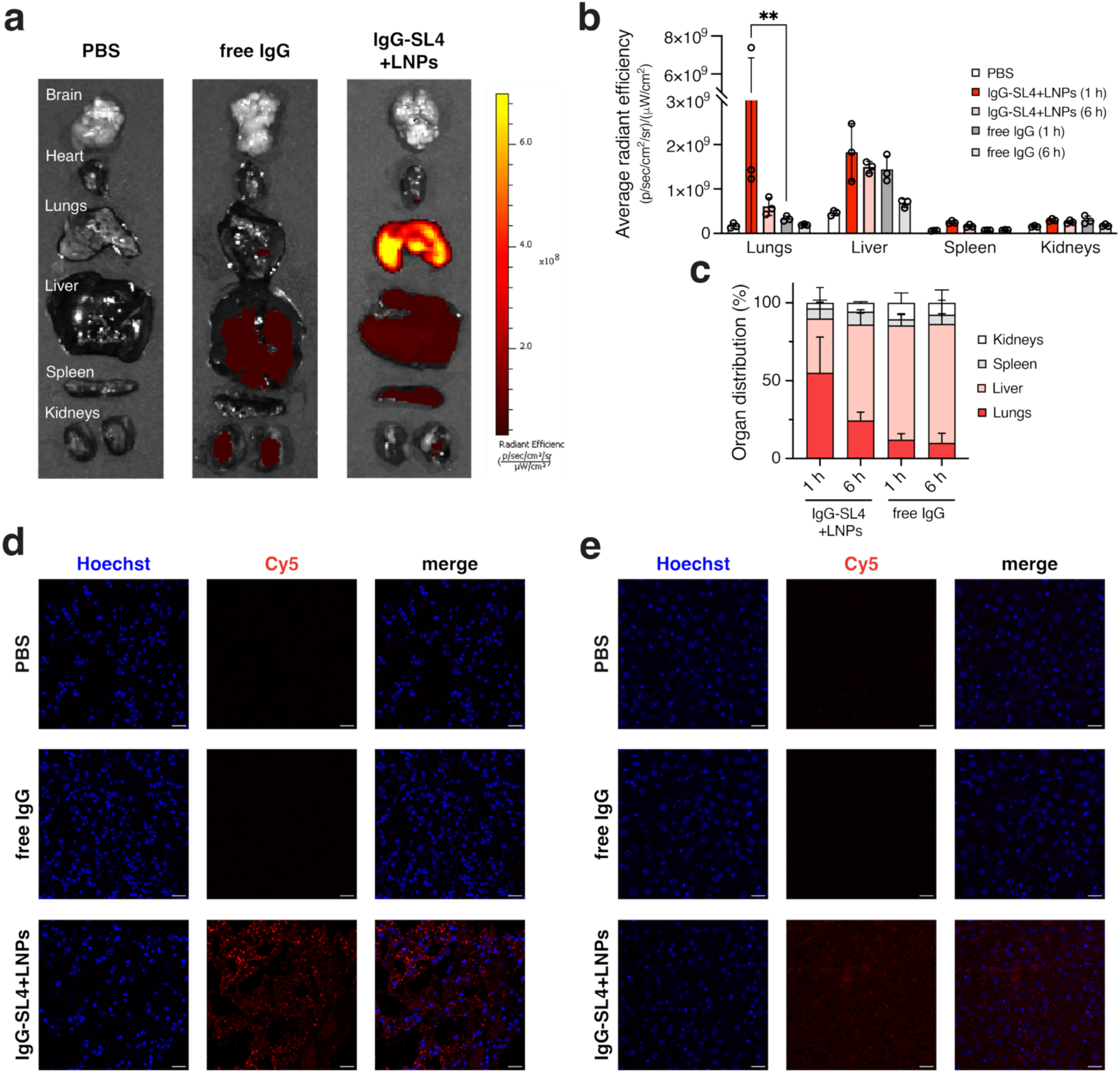
Systemic administration of LNP-encapsulated IgGs results in distribution and uptake in major organs *in vivo*. (a) Representative *ex vivo* fluorescent images of harvested organs following tail vein injections in BALB/c mice of the following: PBS, free IgG, and MC3 LNP-encapsulated IgG. Free and encapsulated IgGs were Cy5-labeled. For encapsulated IgGs, cloaking was performed with 30 molar eq. of SL4 followed by encapsulation in MC3 LNPs (MC3/IgG, 2 wt/wt) supplemented with 30 mol % DOTAP and formulated in PBS pH 5. All injections were performed with 1 mg/kg of total protein. Images shown were taken 1 hr post-injection. (b) Quantified average radiant efficiency of *ex vivo* fluorescent images of harvested lungs, liver, and kidneys from BALB/c mice. (c) Average organ distribution of IgG from harvested lungs, liver, spleen, and kidneys from BALB/c mice. Percentage distribution was calculated by subtracting average PBS radiance from samples. (d-e) Representative confocal microscopy images of sectioned (d) lung and (e) liver tissues of BALB/c mice. Images shown were taken 1 hr post-injection. Scale bar = 20 μm. All data are mean of biological replicates (*n* = 3) ± SD. Statistical significance was determined by two-way ANOVA followed by Bonferroni correction for multiple comparisons (\**p* < 0.05, \*\**p* < 0.01, \*\*\**p* < 0.001, \*\*\*\**p* < 0.0001).

To verify tissue penetration and cellular internalization of IgGs, confocal microscopy was performed on sectioned tissue samples from both the liver and lungs of mice. Notably, a distinct Cy5 signal was observed in the lung and liver tissues of mice administered IgG-encapsulated LNPs, but not in those receiving free IgGs, one hour post injection (**Fig. 4d** and **e**). Furthermore, the Cy5 signal was more pronounced in the lungs compared to the liver, consistent with *ex vivo* fluorescence images of the same organs. Collectively, these findings point towards the general applicability of the cloaking platform to enable LNP-mediated delivery of IgGs to major organs *in vivo*.

### Systemic administration of transferrin-modified LNPs enables targeted delivery of anti-alpha-synuclein antibodies to the brains of a Parkinson’s disease (PD) model

Over the past several years, diseases of the central nervous system (CNS) have emerged as a rising area for therapeutic intervention using LNPs.^38–40^ To extend our established antibody cloaking strategy into neurodegenerative disease models, we investigated the delivery and target engagement of alpha-synuclein (AS)-specific antibodies using SynO4-loaded-LNPs in a Parkinson’s disease (PD) mouse model.^15^ Monoclonal SynO4 antibodies were cloaked, fluorescently labeled, (**Supplementary Figure S13a** and **b**) and subsequently formulated in MC3 LNPs, with a measured size of ∼150 nm and a PDI of 0.153 (**Supplementary Figure S13c**). The LNP encapsulation efficiency was ∼51%, and importantly, the antibody retained activity post-conjugation (**Supplementary Figure S13d** and **e**). *In vitro* delivery confirmed cellular uptake of the cloaked SynO4 with LNPs by flow cytometry (**Supplementary Figure S13f**), validating cellular delivery of the formulation.

We next evaluated whether the system could engage its pathological target *in vivo*. Following intracranial (i.c.) injection of Cy5-SynO4-SL4 LNPs into the substantia nigra (SN) of PD model mice, fluorescent imaging 72 hours post-injection revealed specific colocalization between SynO4 and its target, human phosphorylated alpha-synuclein (pAS) aggregates, in the SN region of the brain (**Fig. 5a–c**). Image analysis revealed a colocalization efficiency of 20.07 ± 9.61%, which suggests localized release and engagement of the antibody *in vivo*. Building upon our previous findings that transferrin-functionalized liposomes improved brain accumulation after systemic administration,^15^ we next engineered transferrin-conjugated LNPs encapsulating Cy5-SynO4-SL4 (**Fig. 5d**). Physical and biochemical analysis confirmed successful LNP fabrication with transferrin conjugation efficiency of ∼70%, a measured size of ∼170 nm, a PDI of 0.095, and a zeta potential of -19.5 mV (**Supplementary Table 2**). Following intravenous (i.v.) injection into PD mice, SynO4 signal was observed in brain regions enriched in pAS, with a colocalization efficiency of 39.30 ± 13.38% (**Fig. 5e** and **f**). These findings demonstrate successful blood–brain barrier (BBB) penetration and target engagement using a non-invasive, systemic route.

**Figure 5.**
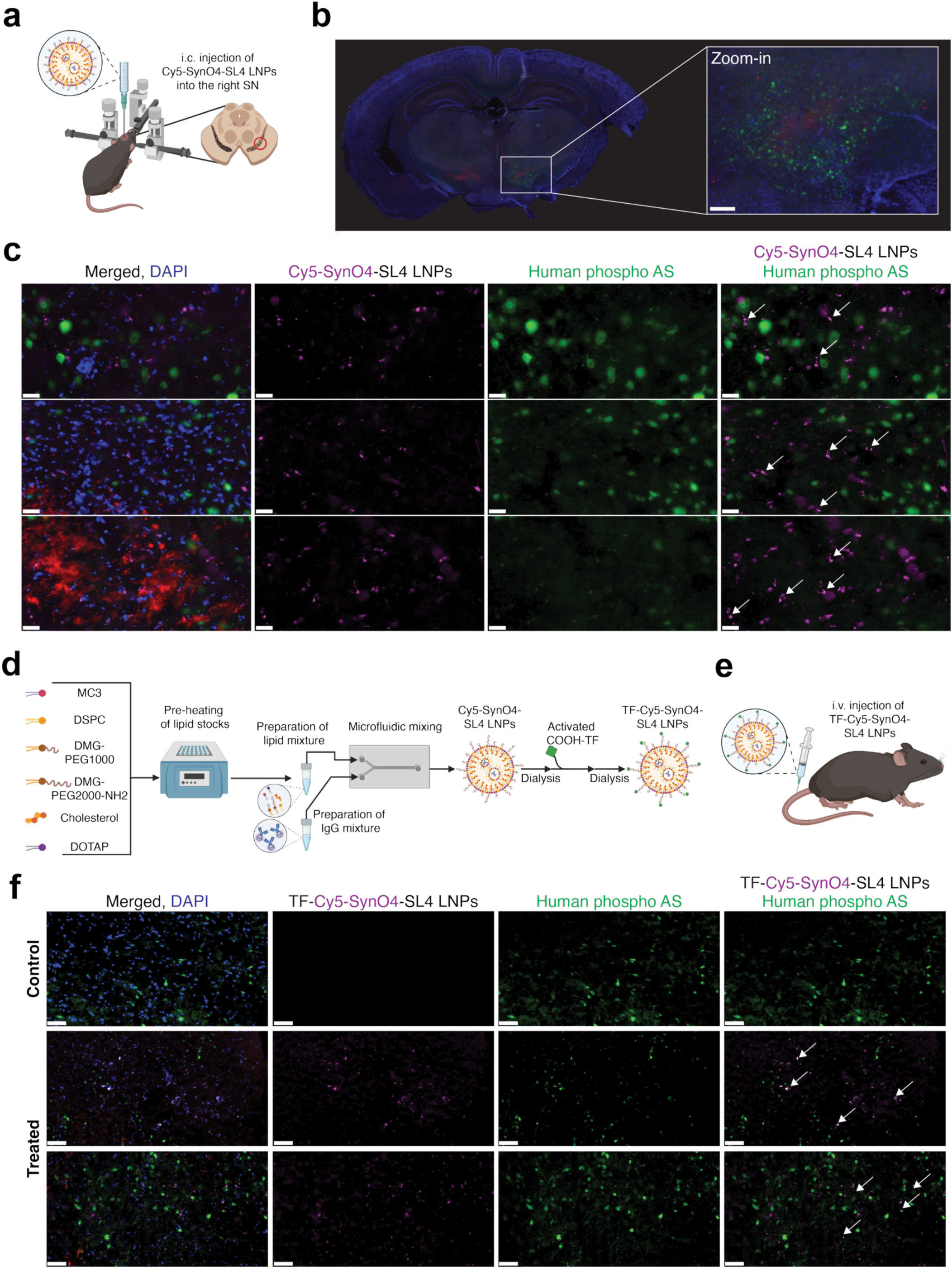
*In vivo* delivery and target engagement of SynO4-loaded LNPs in a Parkinson’s disease (PD) mouse model. (a) Schematic depicting intracranial (i.c.) administration of Cy5-SynO4-SL4-LNPs into the substantia nigra (SN). (b) Representative images of coronal brain tissue sections from the SN region imaged 72 hours after i.c. injection. Sections were fixed and immunostained for tyrosine hydroxylase (TH, dopamine neurons, red), human phosphorylated alpha synuclein (pAS, green), and counterstained with DAPI for cell nuclei (blue). Scale bar = 5 mm. (c) Representative images of brain tissue sections fixed and immunostained for tyrosine hydroxylase (TH, dopamine neurons, red), human phosphorylated alpha synuclein (pAS, green), and counterstained with DAPI for cell nuclei (blue). Colocalization images indicates target engagement of Cy5-SynO4 (purple) with pAS (green). Scale bar = 20 µm. (d) Schematic depicting procedure for formulating transferrin-targeted LNPs (TF-Cy5-SynO4-SL4 LNPs) via microfluidic mixing. (e) Schematic depicting intravenous (i.v.) administration of TF-Cy5-SynO4-SL4 LNPs. (f) Representative images of control brain (non-injected) sections and treated brain sections 6 hours after i.v. injections. Tissue sections were fixed and immunostained for TH (dopamine neurons, red), pAS (green), and counterstained with DAPI for cell nuclei (blue). Cy5-SynO4 antibodies are indicated where they localized and co-localized with pAS in the SN. Scale bar = 50 µm. Results obtained from *n* = 3 independent repetitions (*n* = 9 replicates each).

Together, these results represent a significant advancement in antibody delivery to the CNS using LNP technology. This research expands upon our earlier work with transferrin-liposomes, which increased SynO4 brain accumulation but did not reach levels sufficient to directly visualize antibody colocalization with α-synuclein aggregates.^15^ In contrast, transferrin-LNPs achieved robust parenchymal delivery, enabling clear imaging of SynO4 colocalization with pathological aggregates, and thereby highlight an important advancement in functional target engagement in the brain. Here, the use of LNPs enable enhanced versatility, systemic administration, and transferrin-based active targeting to the brain, laying a foundation for future therapeutic strategies against intracellular protein targets implicated in neurodegenerative diseases.

### Delivery of anti-NF-κB IgGs with lung-targeted LNPs results in attenuation of inflammation in an acute lung injury (ALI) model

Owing to effective signaling inhibition observed *in vitro* and robust biodistribution of IgG-encapsulated LNPs to the lungs, we also proceeded to evaluate their *in vivo* therapeutic efficacy in a mouse model of acute lung injury (ALI). ALI is a clinical pathology that is characterized by extensive lung inflammation and is often observed in infections, sepsis, pneumonia, and severe trauma.^41,42^ ALI is typically associated with tissue damage to the pulmonary endothelium and alveolar epithelium, which can lead to pulmonary edema and hypoxemia (**Fig. 6a**).^41,43^ Recently, it was observed that patients suffering from ALI as a result of severe acute respiratory syndrome coronavirus 2 (SARS-CoV-2) experienced a ∼70% mortality rate,^44^ a debilitating statistic which has motivated increased clinical studies.

**Figure 6.**
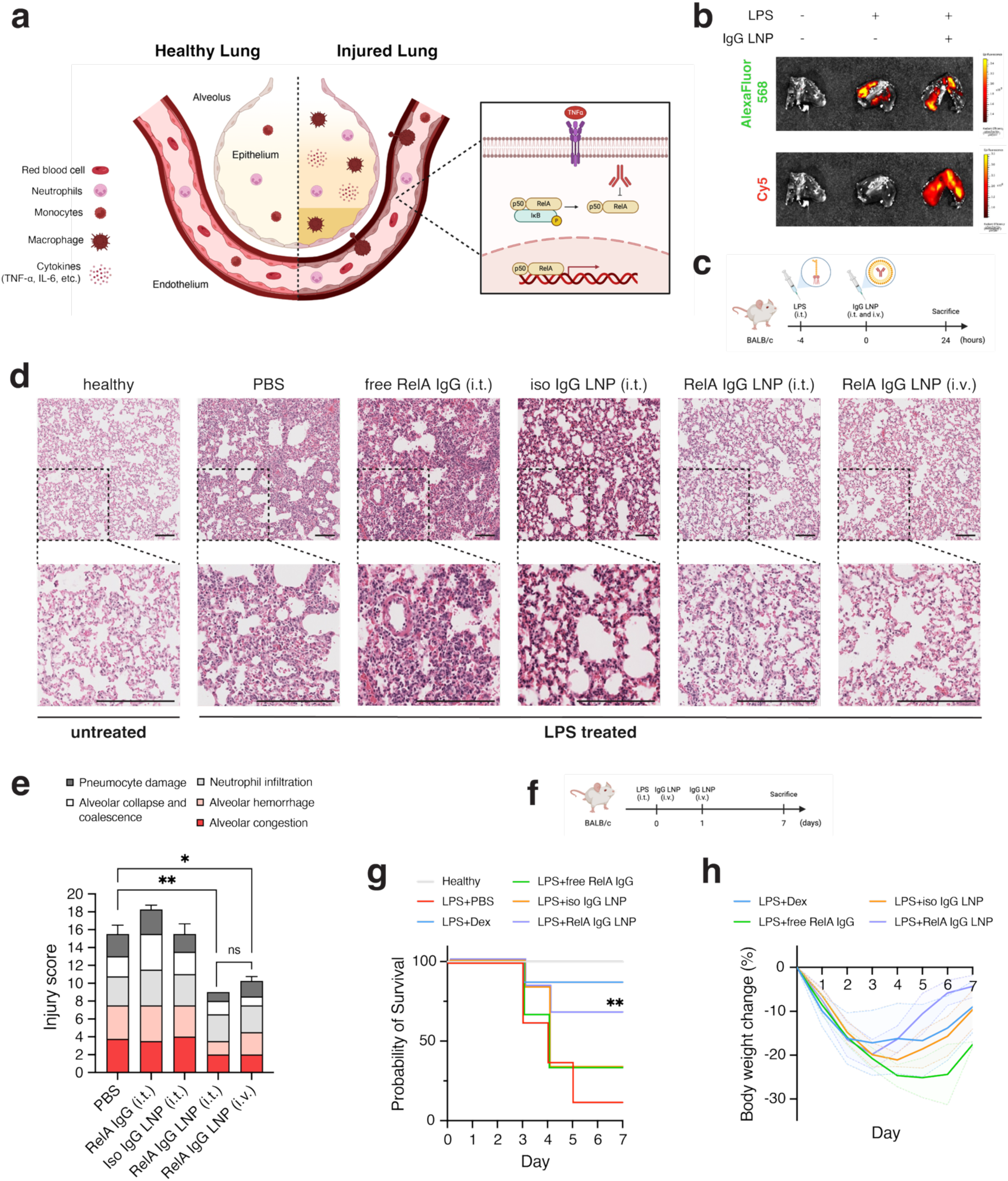
Lung-targeted delivery of LNP-encapsulated anti-NF-κB IgGs leads to reduction of inflammation in ALI mouse model. (a) Schematic depicting lung alveolar damage from inflammation in acute lung injury. (b) *Ex vivo* fluorescent images of harvested lungs following intratracheal injections of AlexaFluor-labeled LPS and Cy5-labeled, MC3 LNP-encapsulated IgGs in BALB/c mice. For encapsulated IgGs, cloaking was performed with 30 molar eq. of SL4 followed by encapsulation in MC3 LNPs (MC3/IgG, 2 wt/wt) supplemented with 30 mol % DOTAP and formulated in PBS pH 5. All injections were performed with 1 mg/kg of total protein. Images shown were taken 1 hr post-injection. (c) Schematic depicting schedule for LPS instillation (3 mg/kg) and LNP-encapsulated IgG administration in BALB/c mice. (d) Histopathologic changes of hematoxylin and eosin-stained lung tissue samples obtained from BALB/c mice following IgG treatments. Scale bar = 100 μm. (e) Semi-quantitative grading of lung tissue damage following IgG treatments. (f) Schematic depicting schedule for LPS instillation and LNP-encapsulated IgG administration in BALB/c mice for longitudinal survival study. (g) Kaplan-Meier survival curve of BALB/c mice following IgG treatments. (h) Weights of survived mice recorded over one week following IgG treatments. All data are presented as mean of biological replicates (*n* = 6–8) ± SD. Statistical significance was determined by one-way ANOVA followed by Bonferroni correction for multiple comparisons and by log-rank test (\**p* < 0.05, \*\**p* < 0.01, \*\*\**p* < 0.001, \*\*\*\**p* < 0.0001).

A hallmark feature of ALI is excessive inflammation caused by pro-inflammatory cytokines such as TNF-α, IL-6, and IL-1β that are produced by infiltrating macrophages in the lung interstitium and alveolus.^45^ In particular, the host inflammatory response to ALI has been well studied and is known to be regulated by the NF-κB pathway.^45,46^ This pathway has emerged as a target for therapeutic interventions aimed at both macrophages and the lung endothelium.^47–49^ Here, we investigated delivering anti-RelA IgGs in lung-targeted MC3 LNPs as a potential therapeutic modality to inhibit ALI-induced inflammation (**Fig. 6a**).

To this end, we first induced ALI in healthy BALB/c mice via intratracheal instillation of lipopolysaccharide (LPS; 3 mg/kg) derived from *Pseudomonas aeruginosa*. Induction of ALI was verified by a loss of body weight as well as an increase in TNF-α levels in the bronchoalveolar lavage fluid (BALF) extracted from LPS-treated mice (**Supplementary Figure S14**). Intratracheal injections of AlexaFluor 568-labeled LPS and Cy5-labeled, cloaked IgGs in LNPs also demonstrated successful localization of LPS and IgG LNPs to the lungs (**Fig. 6b** and **Supplementary Figure S15**), further confirming successful instillation and colocalization of both LPS and IgG LNPs.

Next, we evaluated the therapeutic efficacy of anti-RelA IgGs in ALI-induced mice by analyzing lung tissue damage and measuring changes in pro-inflammatory cytokines (**Figure 6c**). We first assessed the safety of our formulations by examining the lungs of healthy mice administered with LNP-encapsulated isotype control IgG, observing minimal but noticeable inflammation in lung vasculature that can be attributed to the DOTAP-containing formulations (**Supplementary Figure S16**). In ALI-treated mice, histological analysis of sectioned lung tissues obtained 24 hours after administration revealed significantly reduced levels of acute inflammatory response, including attenuated vascular congestion, decreased alveolar immune cell infiltration, and overall reduction of tissue damage to lung endothelium in mice treated with LNP-encapsulated anti-RelA IgG but not in LNP-encapsulated isotype control IgG or free, non-encapsulated anti-RelA IgG (**Figure 6d** and **Supplementary Figure S16**). These observations were further supported by histopathologic assessment by a board-certified anatomic pathologist (R.C.C) with a semi-quantitative grading scale for lung tissue damage (**Figure 6e**). Interestingly, we observed no statistically significant differences in injury scores between intratracheally and intravenously administered LNP-encapsulated anti-RelA IgG (**Figure 6e**). In contrast, analysis of harvested BALF samples by ELISA showed statistically insignificant changes to TNF-α and IL-6 levels (**Supplementary Figure S17**). To reconcile these effects with macrophage inflammatory response, we performed additional experiments using LNP-encapsulated anti-RelA IgGs in RAW264.7 macrophage cells. Here, we observed minimal uptake by flow cytometry (**Supplementary Figure S18**), no reduction in TNF-α levels, and a modest reduction of IL-6 levels in LPS-stimulated RAW264.7 cells treated with RelA IgG LNPs (**Supplementary Figure S19**). Therefore, we theorized that mitigation of the pro-inflammatory M1 macrophage response plays a minimal role in the reduction of lung endothelial damage observed following LNP-encapsulated anti-RelA IgG treatments.

We further extended our efficacy studies in a longitudinal survival model of ALI, where BALB/c mice were instilled with a high dose of LPS (35 mg/kg) followed by two intravenous injections of IgG treatments (1 mg/kg) administered 4 hours and 24 hours after LPS instillation (**Figure 6f**). Mice weights and survival were recorded for a week and compared to those of mice treated with dexamethasone (Dex), an anti-inflammatory drug and widely used ‘gold-standard’. Most ALI-induced mice treated with control samples exhibited low survival rates, with only 10-30% surviving after 5 days (**Figure 6g**). Notably, LNP-encapsulated anti-RelA IgG treatments increased survival to ∼67%, while Dex-treated mice showed over 80% survival (**Figure 6g**). Surviving ALI-induced mice treated with LNP-encapsulated anti-RelA IgG also demonstrated enhanced recovery in weight gain, comparable to that of Dex-treated mice (**Figure 6h**).

Overall, our results showcase the ability of our cloaking platform to deliver anti-inflammatory antibodies in a therapeutic mouse model of ALI. These findings not only highlight the potential for repurposing off-the-shelf antibodies for therapeutic applications but also lay the groundwork for the future development of efficacious antibody treatments targeting intracellular antigens.

## Discussion

Antibodies, considered “magic bullets” in the pharmaceutical industry, are a powerful modality for treating a variety of diseases, owing in large part to their ability to bind to target antigens with remarkably high binding affinity and specificity. In this work, we outline a strategy to introduce antibodies into the cytosol of a cell using cationic lipid vehicles, enabling us to inhibit a variety of different intracellular targets implicated in important disease-associated signaling processes. The ability to deliver antibodies into cells presents significant opportunities to broaden the range of druggable targets within the intracellular environment. This advancement also allows us to leverage well-established pipelines to discover and manufacture antibodies against novel intracellular targets.

Building upon our developed cloaking platform for protein delivery, we demonstrate highly efficient delivery of IgGs into cells using clinically-validated MC3 LNP formulations. We use this strategy to functionally deliver off-the-shelf antibodies against crucial transcription factors, achieving potent inhibition of cell signaling pathways at nanomolar concentrations. These results are especially notable for two reasons. First, many transcription factors have evaded therapeutic intervention due to their “undruggable” structures (as is the case with β-catenin, against which there are no FDA-approved therapies).^50^ Utilizing antibodies, which possess binding paratopes spanning large surface areas, ^51^ could prove to be a compelling alternative to traditional small molecules as a means of inhibiting difficult-to-drug targets. Second, our cloaking platform allows researchers to repurpose a vast market of already existing off-the-shelf antibodies for cell signaling inhibition. By leveraging the platform’s ability to deliver antibodies using both LNP formulations and cationic lipid transfection reagents, we envision this antibody delivery strategy to be incredibly useful for therapeutic development and basic science research of cell signaling pathways.

To investigate potential therapeutic translation, we explored the platform’s ability to deliver IgG cargos across various *in vivo* disease models. Delivery of anti-RelA IgGs using lung-targeted LNPs attenuated inflammation in an ALI model, and transferrin-modified LNPs encapsulating SynO4 antibodies were successfully delivered to the brains of mice with PD. These studies demonstrate the potential of antibodies as efficacious treatments for various diseases and highlight the use of LNP technology to deliver cargos to extrahepatic targets that have traditionally eluded nanoparticle intervention. Ultimately, this work provides a clinically translatable nanotechnology-based platform for delivering antibodies to previously inaccessible intracellular targets, establishing a new framework for therapeutic intervention that could greatly expand the scope and impact of antibody-based medicines.

## Materials and Methods

### Materials

The following chemicals were used as received: Acryloyl chloride (Sigma Aldrich), triethylamine (Acros Organics), propargylamine (Sigma Aldrich), N-Boc-ethylenediamine (Combi-Blocks), 1,8-Diazabicyclo[5.4.0]undec-7-ene (DBU; Sigma Aldrich), sodium 3-mercapto-1-propanesulfonate (3MPS; Sigma Aldrich), 4-mercaptobutyric acid (4MBA; A2B Chem), 2,2-Dimethoxy-2-phenylacetophenone (DMPA; Sigma-Aldrich), trifluoroacetic acid (TFA; BeanTown), 4-nitrophenyl chloroformate (Oakwood), bis(2-hydroxyethyl) disulfide (Sigma Aldrich), and D,L-dithiothreitol (DTT; Sigma Aldrich). DLin-MC3-DMA was purchased from MedChemExpress. Cholesterol was purchased from Sigma Aldrich. 1,2-distearoyl-sn-glycero-3-phosphocholine (DSPC), 1,2-dioleoyl-3-trimethy-lammonium-propane (DOTAP), and 1,2-dimyristoyl-rac-glycero-3-methoxypolyethylene glycol-2000 (PEG-DMG-2000) were purchased from Avanti Polar lipids. Lipofectamine 2000, Lipofectamine 3000, Slide-A-Lyzer MINI Dialysis Devices (3.5 kDa MWCO), Geneticin, Human TNF-α recombinant protein, Human IL-6 recombinant protein, Hoechst 33342, AlexaFluor 488 succinimidyl ester (AlexaFluor 488 NHS ester), and Malachite Green isothiocyanate were purchased from Thermo Fisher Scientific. Micro Float-A-Lyzer Dialysis Devices (1000 kDa MWCO) were purchased from Repligen. Cy5 succinimidyl ester (Cy5 NHS ester) was purchased from BroadPharm. CellTiter 96 Aqueous Non-Radioactive Cell-Proliferation Assay (MTS) and Dual-Glo Luciferase Assay were purchased from Promega. Anti-p50 monoclonal antibody (5D10D11), monoclonal anti-β-catenin, N-terminus antibody (GT2169), anti-β-catenin, C-terminus monoclonal antibody (CAT-5H10), anti-STAT3 monoclonal antibody (9D8), mouse IgG1 isotype control antibody (02-6100), were purchased from Thermo Fisher Scientific. Anti-RelA monoclonal antibody (12H11) and anti-phospho STAT3 monoclonal antibody (9E12) were purchased from Millipore Sigma. NF-κB luciferase reporter plasmid pGL4.32[luc2P/NF-κB-RE/Hygro] and STAT3 luciferase reporter plasmid pGL4.47[luc2P/SIE/Hygro] were purchased from Promega. ELISA Max Deluxe Set Mouse TNF-α and ELISA Max Deluxe Set Mouse IL-6 were purchased from BioLegend.

## Methods

### Chemical synthesis of sulfonated *p*-nitrophenyl carbonate compounds

Compounds were synthesized as done previously (scheme shown in **Supplementary Figure S1**).^28^ Briefly, propargylamine was acylated with acryloyl chloride, followed by a Michael addition with N-Boc-ethylenediamine. Anionic sulfonate groups were installed by reacting the resulting compounds with 3-mercapto-1-propanesulfonate followed by Boc deprotection. Final compounds were purified via RP-HPLC and characterized via mass spectrometry and NMR (**Supplementary Figure S20-21**). For all synthesized materials, NMR spectroscopy was conducted on a Bruker 500 MHz NMR spectrometer. Liquid chromatography-mass spectrometry (LC-MS) was carried out on an Agilent 1200 Series LC/Thermo Fisher LTQ XL MSD equipped with an Agilent EC C18, 2.7 μm, 120 Å LC column (3 x 100 mm, reversed phase), UV diode-array detector monitoring 210 nm, 230 nm, 260 nm, 360 nm, and 505 nm wavelengths, and Agilent multimode source. Masses were detected in either positive or negative ion mode. HPLC purification was performed on a 1100 Series Agilent HPLC system equipped with a UV diode array detector and a 1100 Infinity analytical scale fraction collector using reverse phase C18 column (4.6 x 150 mm, 5 µm).

### Antibody bioconjugation and characterization

For a typical procedure, *p*-nitrophenyl carbonate compounds were dissolved in DMSO to prepare a 10 mg/mL solution. Antibodies were suspended in 10 mM HEPES buffer (pH 8.2) at a concentration of 1 mg/mL and incubated with *p*-nitrophenyl carbonate compounds overnight at 4°C. Conjugated antibody samples were purified by dialyzing against 1X PBS for 24 hours at 4°C to remove unreacted compounds using mini dialysis cups (Slide-A-Lyzer MINI Dialysis Devices, 3.5 kDa MWCO). For encapsulation and cell uptake experiments with fluorophore labeled antibodies, AlexaFluor 488 NHS ester and Malachite Green isothiocyanate were dissolved in DMSO to prepare a 10 mg/mL solution, and the same general reaction procedure was followed. First, antibodies were labeled with NHS ester fluorophores (2 molar eq.) overnight at 4°C, dialyzed for 24 hours at 4°C against 10 mM HEPES buffer (pH 8.2), and then reacted with *p-*nitrophenyl carbonate compounds. Final concentration of conjugated antibody samples was calculated from BCA assay (Thermo Fisher), and conjugation was confirmed by native gel electrophoresis (Bio-Rad) and isoelectric focusing (IEF; Thermo Fisher) as per manufacturers’ protocols.

### Lipid nanoparticle (LNP) formulation and characterization

DLin-MC3-DMA, DSPC, cholesterol, DMG-PEG-2000, and DOTAP were dissolved in ethanol at defined MC3/DSPC/Chol/PEG-2000/DOTAP molar ratios consisting of 50/10/38.5/4.5/11.1 mol/mol for 10 mol% DOTAP formulations and 50/10/38.5/4.5/45 mol/mol for 30 mol% DOTAP formulations (MC3/IgG, 2 wt/wt). Antibody samples were suspended in pH 5 PBS and pipette mixed rapidly into the ethanolic lipid solution at a volume ratio of 3:1 (IgG:lipids, v/v). The resulting mixture was dialyzed (Slide-A-Lyzer MINI Dialysis Devices, 3.5 kDA MWCO) against sterile 1X PBS for 3 hours to remove ethanol. For *in vivo* biodistribution experiments, the formulation mixture was dialyzed (Micro Float-A-Lyzer, 1000 kDa MWCO; Repligen) against sterile 1X PBS overnight at 4°C. Particle size distribution and surface zeta potential of all LNP formulations were measured with a Malvern Zetasizer.

### Encapsulation efficiency

LNP formulations of AlexaFluor 488-labled cloaked antibodies were diluted with 1X PBS to a concentration of 50 μg/mL. 10 μL LNP samples (corresponding to 0.5 μg of total protein) was incubated in the presence or absence of 1% Triton-X (vol/vol), and Triton-X treated samples were sonicated for 5 minutes to lyse LNPs. Samples were mixed with 1 volume equivalent of native gel loading buffer and loaded onto 10% polyacrylamide gels (Bio-Rad) for native gel electrophoresis. Quantification of encapsulation efficiency was calculated from in-gel fluorescence intensities using the following formula: (LNP_lysed_ – LNP_intact_)/LNP_lysed_ × 100. Image analysis and quantification was performed with Bio-Rad Image Lab software.

### Serum stability

LNP formulations of AlexaFluor 488-labled cloaked antibodies were diluted with 1X PBS to a concentration of 50 μg/mL. 10 μL LNP samples (corresponding to 0.5 μg of total protein) was incubated with 10 μL of normal mouse serum (Thermo Fisher) at 37°C for 1 hour. Encapsulation efficiency of proteins following serum incubation was quantified as described above. Serum stability was defined as encapsulation percentage following incubation with serum.

### Cell lines and cell culture

A549, DLD-1, and HepG2 cells were procured from lab stocks. A549 and HepG2 cells were maintained in DMEM media containing 10% FBS and 1% penicillin/streptomycin. DLD-1 cells were maintained in RPMI 1600 media containing 10% FBS and 1% penicillin/streptomycin. All cells were maintained at 37°C, 5% CO_2_ and 90% relative humidity.

### Cell uptake

For a typical antibody delivery procedure, 75,000 cells/well were seeded in a 24-well tissue culture plate and incubated overnight at 37°C. For transfections with Lipofectamine 2000 (Thermo Fisher), 200 nM of antibody samples were diluted in 10 μL of OptiMEM reduced serum media, and 2 μL of Lipofectamine 2000 was suspended in 8 μL of OptiMEM. The protein and lipid reagent solutions were pipette mixed and incubated at room temperature for 10 minutes to allow for complexation. The resulting 20 μL transfection mixture was then added to cells containing 380 μL of cell culture media to a final volume of 400 μL. For transfections with LNPs, LNP solutions formulated with cloaked antibody samples were diluted in 75 μL of 1X PBS and added to 325 μL cell culture media to a final volume of 400 μL. Delivery experiments were performed at 37°C for 6 hours before flow cytometric analysis (see below).

### Flow cytometry

Following delivery experiments in a 24-well tissue culture plate, media was aspirated, and cells were washed three times with PBS. Cells were detached and harvested in 1X PBS by pipetting each well 10-15 times. Cells were pelleted by centrifugation at 500 *g* for 5 minutes, and the supernatant was aspirated and replaced with fresh 1X PBS for flow cytometric analysis. Flow cytometry was performed on an Attune NxT (Thermo Fisher) flow cytometer. FlowJo Version 10 was used to analyze samples by geometric mean fluorescence determined from 10,000 events.

### Luminescence reporter assay

Cells were seeded at 10,000 cells/well in a white-bottom 96-well tissue culture plate and incubated overnight at 37°C. The following day, 100 ng of either TOPFlash plasmid (Addgene), pGL4.32[luc2P/NF-κB-RE/Hygro] (Promega), or pGL4.47[luc2P/SIE/Hygro] (Promega) and 10 ng of Renilla luciferase plasmid (Addgene) was co-transfected into DLD-1, A549, or HepG2 cells, respectively, with Lipofectamine 3000 (Thermo Fisher) in OptiMEM reduced serum media as per manufacturer’s protocol (0.3 μL Lipofectamine 2000/well). 6 hours after transfection, the transfection mixture was replaced with either fresh RPMI 1600 or DMEM media containing 10% FBS and 1% penicillin/streptomycin. MC3 LNPs formulated with cloaked antibodies were then added to each well and incubated for 12 hours 37°C. For NF-κB and STAT3 transcriptional readouts, media was aspirated and replaced with DMEM media containing TNF-α or IL-6 (20 ng/mL) for 5 hours at 37°C. Following incubation, media was aspirated, and cells were washed once with 1X PBS. Cells were lysed, and the firefly and Renilla luminescence signals were measured sequentially by the Dual-Glo Luciferase Assay System (Promega) as per manufacturer’s protocol. Plates were read on a TECAN Infinite M1000 Pro microplate reader. The luciferase activities were measured and normalized against the control Renilla activities.

### Cell viability assay

Cells were seeded at 10,000 cells/well in a in a clear 96-well tissue plate and incubated overnight at 37°C for delivery experiments. After experiment, media was aspirated, and cells were washed once with 1X PBS. 100 μL of FluoroBrite DMEM and 10 μL of MTS solution (Promega) were added to each well, and the plate was incubated for 1 hour. Absorbance measurements were taken at 490 nm on a TECAN Infinite M1000 Pro microplate reader and normalized to untreated cells (100%) or clear media (0%).

### Fluorogen activating protein (FAP) assay

To generate dL5-mCer3 expressing cell lines, A549 cells were first seeded at a density of 300,000 cells/well in a 6-well tissue culture plate and incubated overnight at 37°C. The following day, cells were transfected with 2.5 μg of pcDNA3.1-KozATG-dL5-2XG4S-mCer3 (Addgene) with Lipofectamine 3000 (Thermo Fisher) in OptiMEM reduced serum media as per manufacturer’s protocol (2.5 μL Lipofectamine 3000/well). 6 hours after transfection, the transfection mixture was replaced with fresh DMEM media containing 10% FBS and 1% penicillin/streptomycin. The following day, cells were harvested and reseeded at a density of 150,000 cells/well in a 6-well tissue culture plate with media containing Geneticin (200 μg/mL) to select for transfected cells. Once cells reached 70-80% confluency, cells were harvested and reseeded at a density of either 75,000 cells/well in a 24-well tissue culture plate (for FACS) or 10,000 cells/cm^2^ in a 35 mm^2^ cell culture dish (for microscopy) and incubated overnight at 37°C. MC3 LNPs formulated with mGe-labeled antibody samples were then added to cells and incubated for 6 hours at 37°C before analysis via FACS and confocal microscopy.

### Confocal microscopy

Cells were seeded at 10,000 cells/cm^2^ in a 35 mm^2^ cell culture mm^2^ dish and incubated overnight at 37°C. MC3 LNPs formulated with formulated with mGe-labeled antibody samples were added to each dish in a total volume of 2 mL of cell culture media. Delivery experiments were performed at 37°C for 6 h. Media was aspirated, and cells were washed three times with 1X PBS. 2 mL of FluoroBrite DMEM media containing SYTO green diluted 1:20,000 was added to each dish. After 10 minutes of incubation, media was aspirated, and cells were washed three times with 1X PBS. Dishes were then refilled with 2 mL of FluoroBrite DMEM before imaging. Samples were imaged on an inverted Zeiss LSM88-confocal microscope (i880) using a 40×water immersion objective. Images were analyzed with FIJI software.

### Enzyme-linked immunosorbent assay (ELISA) of anti-RelA IgGs against RelA

A 96-well enzyme immunoassay plate was coated with 100 μL of recombinant RelA protein (Novus Biologics NBP2249875UG) at 4 μg/mL in 0.05 M NaCO3 buffer (pH 9.6) overnight at 4°C. The plate was washed three times with 200 μL of PBST (1X PBS and 0.1% Tween) per well and blocked with 200 μL of 1X PBS containing 3% milk per well overnight at 4°C. The plate was washed three times with 200 μL of PBST per well. Serial dilutions of anti-RelA antibody samples in blocking buffer were added at 50 μL per well, and the plate was slowly mixed for 1 hour at room temperature. The plate was washed three times with 200 μL of PBST per well and then incubated with anti-mouse antibody conjugated to horsereadish peroxidase (HRP) diluted 1:10,000 in 50 μL PBST+1% milk for 1 hour with slow mixing. The plate was washed three times with 200 μL of PBST per well before the addition of 50 μL of 1-Step Ultra TMB (Thermo Fisher). The reaction was allowed to incubate with slow mixing and then quenched with 50 μL of 3N H_2_SO_4_. Absorbance measurements were taken at 450 nm on a TECAN Infinite M1000 Pro microplate reader.

### Co-immunoprecipitation (co-IP) of RelA and Western Blot analysis

A549 cells were seeded at 300,000 cells/well in a 6-well tissue culture plate and incubated overnight at 37°C. The following day, 200 nM of MC3 LNPs formulated with cloaked isotype IgG3 antibodies and anti-RelA antibodies were then added to each well and incubated for 12 hours at 37°C. Media was aspirated and replaced with DMEM media containing TNF-α (20 ng/mL) for 5 hours at 37°C. Following incubation, media was aspirated, and cells were washed once with 1X PBS. Cells were lysed on ice for 10 minutes with 100 μL of Pierce RIPA lysis buffer premixed with 1X Halt protease inhibitor and 1X EDTA solution and then pelleted by centrifugation at 15,000 *g* for 15 minutes. The supernatant was collected, and total protein content was quantified using BCA assay (Thermo Fisher). 50 μg of cell lysate was then diluted to 50 μL of 1X PBS and incubated with 50 μL of Dynabeads™ Protein G (Thermo Fisher) on a rotator overnight at 4°C. Samples were washed three times with 1X PBS, resuspended in 1X PBS, and boiled for subsequent Western Blot analysis. 25 μg of lysates were separated on Any kD polyacrylamide gels (Bio-Rad), transferred to nitrocellulose membranes, and blocked overnight with 1X TBS containing 5% milk. Membranes were then incubated with mouse anti-RelA antibody (Millipore Sigma 12H11) diluted 1:1,000 in TBST (1X TBS and 0.1% Tween)+1% milk for 1 hour at room temperature. Membranes were washed with 1X TBST and then incubated with secondary anti-mouse antibody conjugated to HRP diluted 1:10,000 in TBST +1% milk for 1 hour at room temperature. Membranes were washed with 1X TBST before imaging.

### TRIM21 silencing reporter assay

A549 cells were seeded at 10,000 cells/well in a white-bottom 96-well tissue culture plate and incubated overnight at 37°C. The following day, 100 ng of pGL4.32[luc2P/NF-κB-RE/Hygro] (Promega) and 10 ng of Renilla luciferase plasmid (Addgene) was co-transfected with Lipofectamine 3000 (Thermo Fisher) in OptiMEM reduced serum media as per manufacturer’s protocol (0.3 μL Lipofectamine 2000/well). 6 hours after transfection, the transfection mixture was replaced with fresh DMEM media containing 10% FBS and 1% penicillin/streptomycin. 100 nM of cloaked RelA antibodies formulated with MC3 LNPs as well as 100 ng of either human TRIM21 siRNA (Thermo AM16708) or negative control siRNA (Thermo 4390843) formulated with MC3 LNPs (2 wt/wt, MC3/siRNA) were then added to each well and incubated for 12 hours 37°C. Media was aspirated and replaced with DMEM media containing serial dilutions of TNF-α (0.1-100 ng/mL) for 5 hours at 37°C. Following incubation, media was aspirated, and cells were washed once with 1X PBS. Cells were lysed, and the firefly and Renilla luminescence signals were measured sequentially by the Dual-Glo Luciferase Assay System (Promega) as per manufacturer’s protocol. Plates were read on a TECAN Infinite M1000 Pro microplate reader. The luciferase activities were measured and normalized against the control Renilla activities.

### *In vivo* biodistribution

Female BALB/c mice (strain # 000651, 6–7 weeks old) were obtained from Jackson Laboratory. All mice were injected intravenously (i.v.) through the tail-vein. Before injections, total protein amount of MC3 LNPs formulated with Cy5-labeled antibodies was calculated from BCA assay of LNP samples. Mice weighing 18-20 g were i.v. injected with 1X PBS as well as free antibodies and MC3 LNPs formulated with Cy5-labeled cloaked antibodies at a dose of 1 mg/kg of total protein. *Ex vivo* imaging of harvested organs was performed 1 hour and 6 hour post injection on an IVIS Spectrum system (Perkin Elmer). Images were analyzed with Living Image (Perkin Elmer) software. All animal experiments were reviewed and approved by the Institution of Animal Care and Use Committees (IACUC) of Cornell University under protocol # 2019-0063.

### ELISA of SynO4 IgGs against α-synuclein

An in-house ELISA assay using the direct ELISA method was used for the Anti-Human SNCA Therapeutic (SynO4) Antibody (TAB-0750CLV-L; Creative Biolabs, USA). A 384-well plate was coated with 1.4 µg/mL of recombinant human alpha-synuclein protein aggregate (Active) (ab218819; Abcam, UK), sealed, and incubated overnight at 4 °C. The next day, plates were blocked in 5% fetal bovine serum (FBS) (K210430; Rhenium, Israel) in PBST (Dulbecco’s phosphate-buffered saline [PBS]) (D8537; Sigma-Aldrich, Israel) with 0.05% Tween20 (P1379; Sigma-Aldrich); this was followed by the addition of known concentrations of SynO4 antibody and by the addition of different SynO4 forms samples: SynO4, SynO4-Cy5, SynO4-SL4-Cy5, SynO4-SL4-Cy5 post treatment with DTT. Next, plates were washed to remove PBST, followed by incubation with the secondary antibody (1:100,00; Goat Polyclonal to Mouse IgG1-HRP) (ab6789; Abcam). TMB ELISA Substrate (ab171522; Abcam) was used for signal development, with kinetics absorbance measured using a plate reader (reading at 650 nm with 2 min intervals for 1 hour). A linear calibration curve for the SynO4 antibody was generated based on the described ELISA setup, and a graph of the antibody activity percentage was generated (**Figure S13e**).

### Transferrin-modified LNP formulation and characterization

A total lipid mixture of 4-(dimethylamino)-butanoic acid, (10Z,13Z)-1-(9Z,12Z)-9,12-octadecadien-1-yl-10,13-nonadecadien-1-yl ester (DLin-MC3-DMA), (34364; Cayman, USA), 1,2-Distearoyl-sn-glycero-3-phosphoethanolamine (DSPE), (565400; Lipoid, Germany), cholesterol (C8667; Sigma-Aldrich), 1,2-Dimyristoyl-sn-glycero-3-methoxypolyethylene glycol l000 (DMG-PEG1000), (001317-1K; Biopharma PEG, USA), 1,2-dimyristoyl-rac-glycero-3-polyethylene glycol 2000 amine (DMG-PEG2000-NH2), (DMG-PEG2k-AM; NANOCS, USA), and 1,2-dioleoyl-3-trimethylammonium-propane (chloride salt) (DOTAP), (593510; Lipoid, Germany) in molar percentages of 34.0:6.8:26.2:2.4:0.5:30, was dissolved in ethanol (the organic phase). Cy5-SynO4-SL4 was dissolved in PBS (pH 5) to produce an aqueous phase. Using the NanoAssemblerIgnite (NIN0001; Cytiva, USA; provided by A. Zinger lab, Technion), a microfluidic device, the organic and aqueous phases were combined at a 1:5 volumetric ratio and flow rate of 12 ml/min and diluted in a 1:1 volume ratio in PBS to produce LNPs. LNPs were then dialyzed against PBS (pH 7.4; 1:1000 volume ratio) using a 1000 kDa dialysis membrane (131486; Repligen, USA) to change the organic solvent to water-based buffer and to remove the unencapsulated Cy5-SynO4-SL4 antibodies at 4°C for 1 hour. Human holo-transferrin (T4132; Sigma-Aldrich) was cross-linked to the surface of Cy5-SynO4-SL4 LNPs using N-(3-dimethylaminopropyl)-N’-ethyl carbodiimide hydrochloride (EDC) (03450; Sigma-Aldrich) and N-hydroxysulfosuccinimide sodium salt (Sulfo-NHS) (FH24507; Tzamal D-Chem Laboratories Ltd, Israel). First, 10 mg/mL of TF (PBS), 15 mg/mL of Sulfo-NHS (PBS), and 15 mg/mL of EDC (DMSO) solutions were prepared. Second, each reagent solution was added to the TF solution to reach a 1:25 molar proportion (TF:Sulfo-NHS) and 1:10 molar proportion (TF:EDC). Using 1.2 M HCl, the pH of the TF solution with reagents was adjusted to 6. Third, the solution was mixed (450 rpm) at 25 °C for 1 hour. Fourth, the TF solution was dialyzed against PBS (pH 6; 1:1000 volume ratio) using a 12–14 kDa dialysis membrane (132700; Repligen) at 4°C overnight to remove excess reagents. Fifth, TF-Cy5-SynO4-SL4 LNPs were synthesized by mixing Cy5-SynO4-SL4-LNPs with the TF solution in a molar proportion of 10:1 (LNPs:TF). Sixth, the pH of the LNP reaction was adjusted to 8.4 with 1 M sodium bicarbonate, and the reaction was incubated for 2 hours at 450 rpm and maintained at 25 °C. Last, non-conjugated TF was removed by dialysis against PBS (pH 7.4; 1:1000 volume ratio) using a 1000 kDa dialysis at 4°C overnight. To monitor progress, samples were measured using the Zetasizer Ultra (Malvern) after each dialysis step (**Supplementary Table 2**). Micro BCA Protein Assay Kit was used to quantify TF conjugation efficiency (**Supplementary Table 2**) on the surface of the LNPs.

### Adenovirus (AAV)-based Parkinson’s disease (PD) mouse model

Male C57BL/6JRccHsd mice (6–8 weeks old; Envigo, Israel) were anesthetized with 0.5% isoflurane in 1% O₂ and placed in a stereotactic frame. Mice received a unilateral injection of 1.5 µL recombinant AAV2/6 encoding human SNCA (rAAV2/6-hSyn1-Human SNCA-WPREpolyA; 1.63 × 10¹³ GC/mL; Sirion Biotech, Germany) into the right substantia nigra using an automated stereotactic injector (coordinates based on the mouse brain atlas). Mice were monitored daily for 4 weeks. At 4 weeks post-injection, mice were administered either 350 µL of TF-Cy5-SynO4-SL4-LNPs (100 µg/mL SynO4) via tail vein injection or 2 µL of Cy5-SynO4-SL4-LNPs (100 µg/mL SynO4) directly into the right substantia nigra via stereotactic injection. Mice receiving intravenous injections were sacrificed after 6 hours; those receiving stereotactic injections were sacrificed after 72 hours. Euthanasia was performed via Ketamine/Xylazine overdose, followed by transcardial perfusion with PBS. Brains were dissected, fixed in 4% paraformaldehyde at 4°C overnight, and cryoprotected in 15% and 30% sucrose (in PBS) for 24 hours.

### Floating sectioning and immunohistochemistry

PD mice brains were embedded in OCT and cryosectioned at 50 μm into free-floating coronal sections throughout the full rostrocaudal extent. Sections were permeabilized in 0.3% Triton X-100 in PBS for 5 min at room temperature, followed by blocking in 10% normal goat serum (NGS; S-1000; Vector Laboratories Inc, US) for 1 hour. Primary antibodies were incubated overnight at 4 °C in blocking buffer: chicken anti-tyrosine hydroxylase (1:1,000; ab76442, Abcam) and rabbit anti-alpha-synuclein (1:2,000; ab178846, Abcam). After washing, sections were incubated for 1 hour at room temperature with secondary antibodies diluted in blocking buffer: Alexa Fluor 488-conjugated goat anti-rabbit IgG (1:250; ab150077, Abcam) and Alexa Fluor 568-conjugated goat anti-chicken IgY (1:1,000; ab175711, Abcam). Sections were counterstained and mounted using DAPI Fluoromount-G (010020; ENCO, Israel), cover-slipped, and dried overnight at 4°C. Slides were imaged using a 3DHISTECH Pannoramic 250 Flash III automated slide scanner.

### Fluorescence image quantification and overlay

Fluorescence microscopy images for Cy5 and FITC channels were analyzed in batches using Python 3.12.4. For each field, grayscale images from both channels were subjected to fixed thresholding (24 for Cy5, 12 for FITC) to generate binary masks of positive signal. The area of each mask and their overlap were quantified by counting positive pixels. For visualization, color overlays were generated: FITC-positive regions were shown in green, Cy5-positive regions were shown in magenta, and where both signals overlapped, the region appeared white to indicate simultaneous presence of both markers. All masks, merged images, and corresponding quantification data were saved in excel file for further analysis.

### Acute lung injury (ALI) mouse model

Female BALB/c mice (strain # 000651, 6–7 weeks old) were obtained from Jackson Laboratory. To induce lung injury, mice were anaesthetized then intratracheally (i.t.) administered LPS (2 mg/mL in 1X PBS) at a dose of 3 mg/kg via using a 22-gauge catheter. 4 hours after LPS instillation, 1X PBS as well as free antibodies and MC3 LNPs formulated with cloaked antibodies were injected into mice at a dose of 1 mg/kg via i.t. injection and/or i.v. injection through the tail-vein. 24 hours after LNP injections, bronchoalveolar lavage fluid (BALF) from mice were extracted by instillation and extraction with 600 μL of 1X PBS. Collected BALF samples were centrifuged at 400 *g* for 5 minutes at 4°C, and the supernatant was collected for subsequent analysis. After euthanasia, lung tissues were also harvested from all mice for subsequent histopathological analysis.

For survival model of ALI, mice were administered LPS (15 mg/mL in 1X PBS) at a dose of 35 mg/kg, and 4 hours later, 1X PBS as well as free antibodies and MC3 LNPs formulated with cloaked antibodies were injected into mice at a dose of 1 mg/kg via i.v. injection through the tail-vein. Dexamethasone-treated mice were administered dexamethasone (1 mg/mL in 5% DMSO in water) at a dose of 1 mg/kg via intraperitoneal injection (i.p.) 30 minutes before LPS instillation. 24 hours later, a second set of i.v. injections were again performed with the same PBS and antibody samples. Mice weights and survival was recorded daily for one week followed by euthanasia. All animal experiments were reviewed and approved by the Institution of Animal Care and Use Committees (IACUC) of Cornell University under protocol # 2019-0063.

### ELISA of pro-inflammatory cytokines from BALF samples

Levels of the inflammatory cytokines interleukin-6 (IL-6) and tumor necrosis factor-α (TNF-α) were determined by enzyme-linked immunosorbent assay (ELISA) as per manufacturer’s protocol (BioLegend). Briefly, 96-well enzyme immunoassay plate was coated with supplied capture antibody in the supplied coating buffer overnight at 4°C. The plate was washed four times with 200 μL of PBST (1X PBS and 0.1% Tween) per well and blocked with 200 μL per well of supplied blocking buffer for 1 hour at room temperature. The plate was washed four times with 200 μL of PBST per well. BALF samples diluted in supplied blocking buffer were added at 100 μL per well, and the plate was slowly mixed for 2 hours at room temperature. The plate was washed four times with 200 μL of PBST per well and then incubated with 100 μL of supplied detection antibody solution for 1 hour with slow mixing. The plate was washed four times with 200 μL of PBST per well and then incubated with 100 μL of supplied Avidin-HRP solution for 30 minutes with slow mixing. The plate was washed five times with 200 μL of PBST per well before the addition of 100 μL of supplied TMB substrate. The reaction was allowed to incubate with slow mixing and then quenched with 100 μL of supplied stop buffer. Absorbance measurements were taken at 450 nm on a TECAN Infinite M1000 Pro microplate reader.

### Histology

Biopsy punches measuring 4 mm in diameter were taken from harvested lung and liver samples of BALB/c mice and placed in histology cassettes. Samples were fixed in 10% neutral buffered formalin for 48 hours and then placed into 1X PBS for 24 hours. Fixed tissue samples were trimmed and paraffin embedded for routine histology processing by the Animal Health Diagnostic Center at the Cornell University College of Veterinary Medicine. For confocal microscopy, histology slides were incubated with PBS containing Hoechst diluted 1:10,000 for 10 minutes before imaging. Samples were imaged on an inverted Zeiss LSM88-confocal microscope (i880) using a 40X water immersion objective. Images were analyzed with FIJI software.

### ELISA of pro-inflammatory cytokines from RAW264.7 cells

A total of 10,000 RAW264.7 cells per well were plated in a clear 96-well tissue plate and incubated overnight at 37°C. The following day, LPS (100 ng/well) was added to each well for 3 hours. Following incubation, media was aspirated, and cells were washed once with 1X PBS and replaced with fresh DMEM media. Cloaked RelA antibodies formulated with MC3 was then added to each well and incubated for 24 hours 37°C. 90 μL of media was collected and diluted 3X with dilution assay buffer and subsequently analyzed by ELISA as per manufacturer’s protocol (BioLegend), described previously.

