## Supplemental Data for "Intracellular delivery of full-length antibodies via organ targeted lipid nanoparticles"

### Supporting Information

| | Z-average (nm) | PDI | Zeta potential (mV) |
|---|---|---|---|
| **IgG-SL4+LNPs** | 223 ± 6.3 | 0.14 ± 0.016 | -0.48 ± 0.26 |

**Supplementary Table 1. Dynamic light scattering of MC3 LNPs formulated via manual mixing with cloaked isotype IgGs.** Size (z-avg), PDI, surface zeta potential of MC3 LNPs (MC3/IgG, 2 wt/wt) formulated with IgG-SL4 (modified with 30 molar eq.). LNPs were formulated in pH 5 PBS supplemented with 30 mol% DOTAP. All data are mean ± SD (*n* = 3).

| | Z-average (nm) | PDI | Zeta potential (mV) | Total number conc. (particles/ml) | Transferrin conjugation efficiency (%) |
|---|---|---|---|---|---|
| **Post-microfluidic mixing** | 104.6 ± 6.5 | 0.1594 ± 0.0493 | - | - | - |
| **Post-first dialysis** | 114.2 ± 2.3 | 0.1365 ± 0.0327 | - | - | - |
| **Post-transferrin conjugation** | 141.5 ± 11.3 | 0.1102 ± 0.0425 | - | - | - |
| **Post-final dialysis** | 170.1 ± 17.3 | 0.0946 ± 0.0212 | -19.54 ± 1.11 | $3.13 \times 10^{10}$ ± $0.43 \times 10^{10}$ | 69.5 ± 29.0 |

**Supplementary Table 2. Dynamic light scattering and characterization of MC3 LNPs formulated with cloaked SynO4 IgGs.** Size (z-avg), PDI, surface zeta potential, and particle concentration of MC3 LNPs formulated with SynO4 IgG-SL4 (modified with 30 molar eq.). LNPs were formulated in pH 5 PBS supplemented with 30 mol% DOTAP. Transferrin conjugation efficiency was quantified by BCA assay. All data are mean ± SD (*n* = 3).

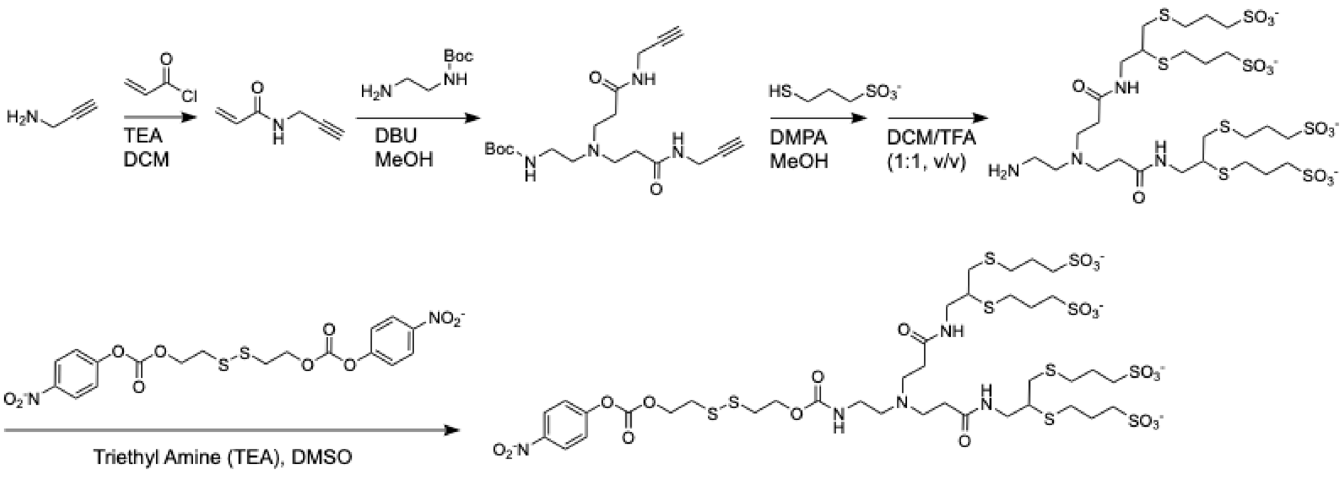


**Supplementary Figure S1. Synthesis scheme for cloaking reagent.** Scheme outlining general synthetic steps for producing SL4.

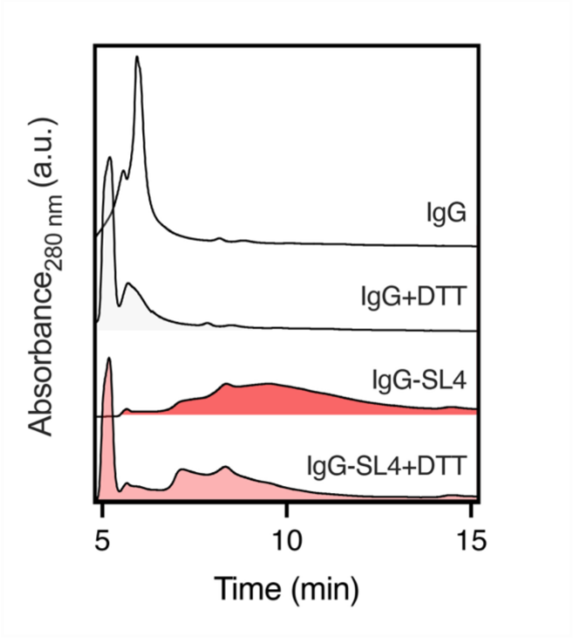


**Supplementary Figure S2. Anion exchange chromatography of DTT-treated IgG samples.** Chromatograms (measured at 280 nm) from anion exchange chromatography of IgG samples, before and after treatment with 10 mM DTT. IgGs were cloaked with 30 molar eq. of SL4.

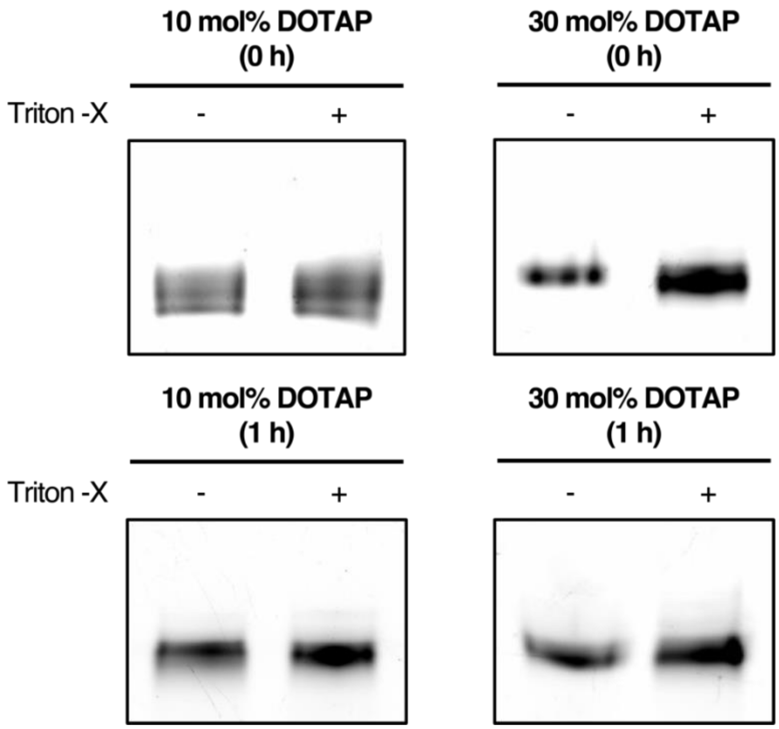


**Supplementary Figure S3. Encapsulation efficiency of cloaked IgGs formulated with LNPs.** Representative in-gel fluorescence image of gel electrophoresis run under native conditions of IgG-SL4 (modified with 2 molar eq. of NHS-AlexaFluor 488 and 30 molar eq. of SL4) formulated with MC3 LNPs, before and after incubation with mouse serum for 1 hour at 37°C. Data shown are for MC3 LNPs (MC3/IgG, 2 wt/wt) supplemented with 10 mol% and 30 mol % DOTAP and formulated in pH 5 PBS. LNP samples were treated with Triton-X to dissolve LNPs and release encapsulated proteins. Gels were loaded with 0.5 µg protein/well. Percent encapsulation was calculated by normalizing signal from LNP samples with signal from LNP samples treated with Triton-X.

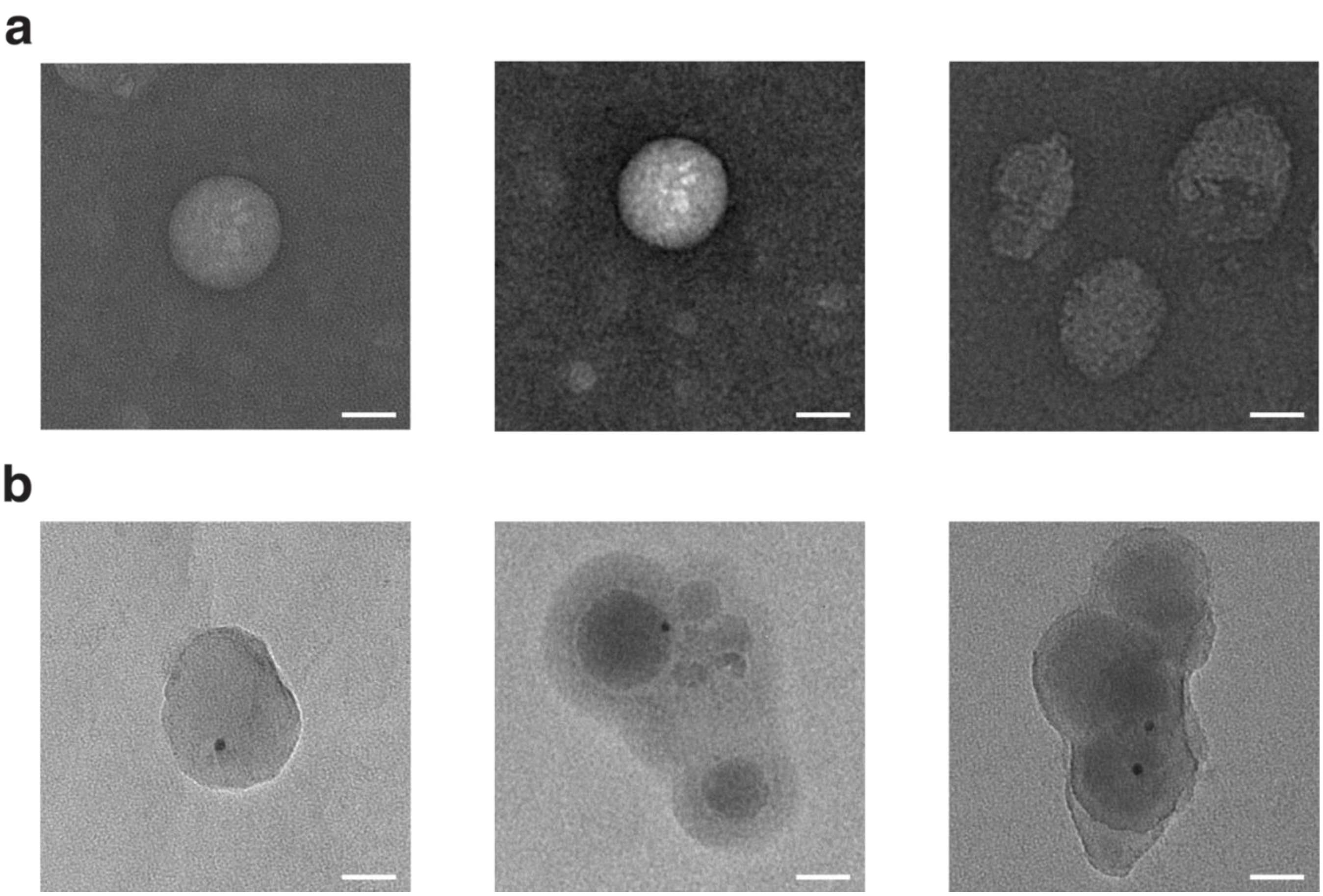


**Supplementary Figure S4. TEM images of cloaked IgGs formulated with LNPs.** (a) Representative TEM images of MC3 LNPs (MC3/IgG, 2 wt/wt) formulated with IgG-SL4 (modified with 30 molar eq. of SL4). (b) Representative TEM images of MC3 LNPs (MC3/IgG, 2 wt/wt) formulated with NanoGold® IgG-SL4 (modified with 30 molar eq. of SL4). LNPs were formulated in pH 5 PBS supplemented with 30 mol% DOTAP. Scale bar = 100 nm.

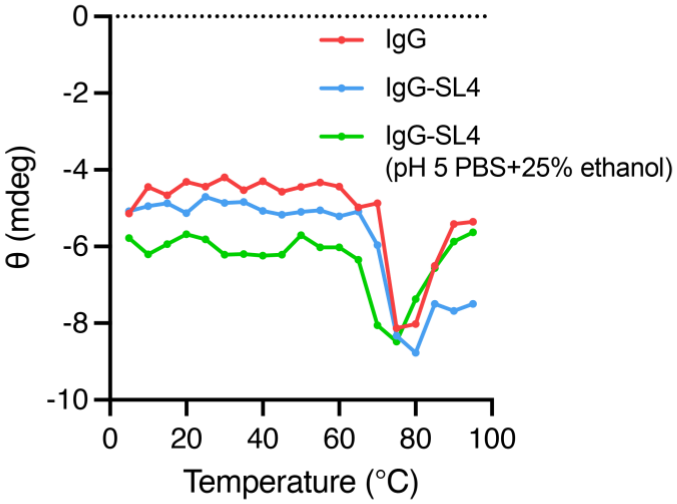


**Supplementary Figure S5. Structural analysis of cloaked IgG samples.** CD spectra of native IgG and cloaked IgG samples. Cloaked IgG samples were placed in pH 5 buffer containing 25% ethanol to mimic formulation conditions (green). Scans were performed at 218 nm using a temperature scan of 5°C-95°C with a 5°C interval.

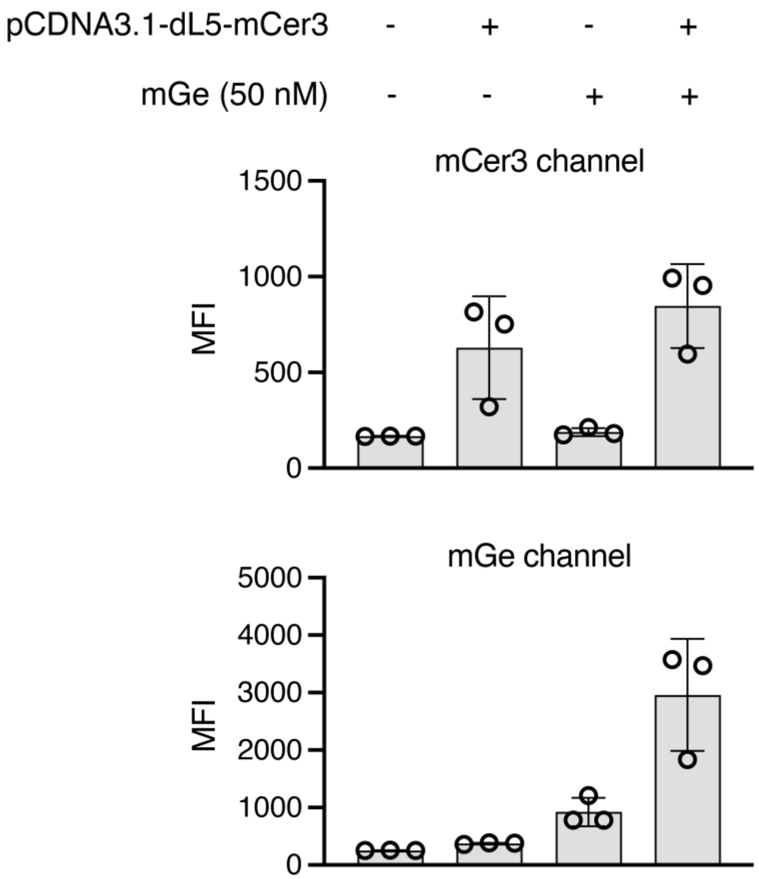


**Supplementary Figure S6. Flow cytometry results of A549 cells using fluorogen activating protein assay.** Mean fluorescence intensity of cells after transfection of pCDNA3.1-dL5-mCer3 followed by 6 hour incubation with malachite green isothiocyanate (50 nM). Fluorescence readings were obtained in channels for mCer3 and mGe. All data are presented as mean ± SD (*n* = 3 for flow cytometry).

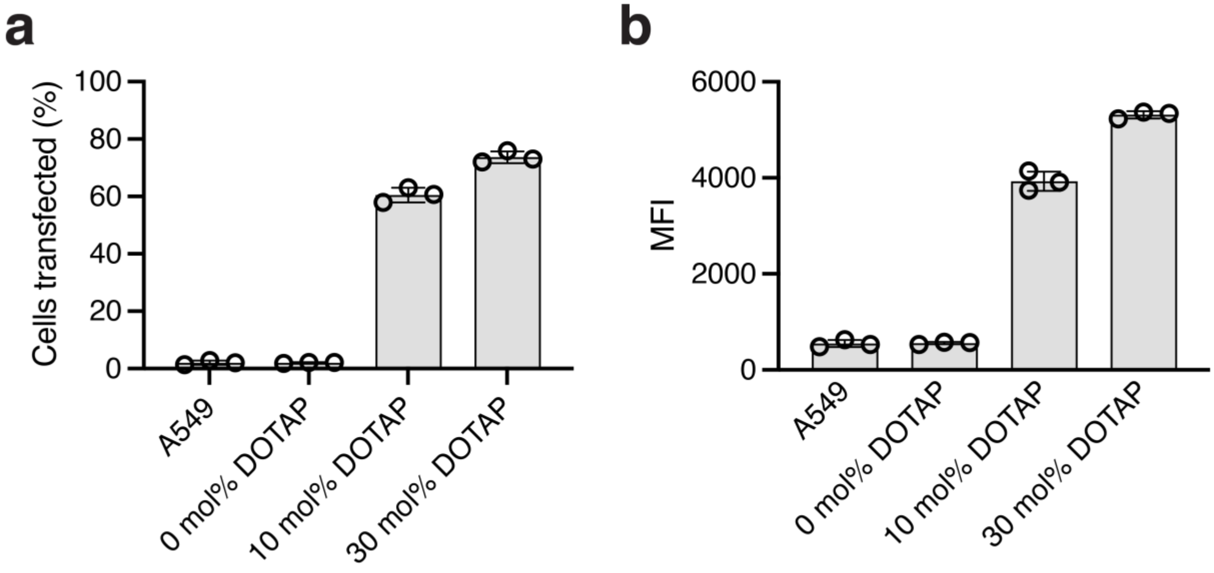


**Supplementary Figure S7. Flow cytometry results of A549 cells treated with IgG-encapsulated LNPs using fluorogen activating protein assay.** Data shown are for IgGs cloaked with 30 molar eq. of SL4 and for MC3 LNPs (MC3/IgG, 2 wt/wt) supplemented with 0-30 mol % DOTAP and formulated in pH 5 PBS. All transfections performed for 6 hours. (a) Percent positive mGe labeled IgG-positive cells following transfections of cloaked IgGs using MC3 LNPs. (b) Mean fluorescence intensity of cells (obtained in mGe channel) following transfections of cloaked IgGs using MC3 LNPs. All data are presented as mean ± SD (*n* = 3 for flow cytometry).

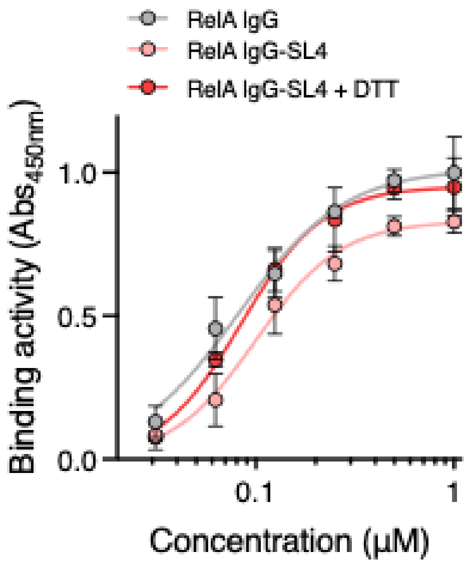


**Supplementary Figure S8. ELISA of anti-RelA IgG against RelA.** Binding activity of anti-RelA IgG to immobilized RelA as determined by ELISA in the presence or absence of DTT. All data are presented as mean ± SD (*n* = 3 for ELISA).

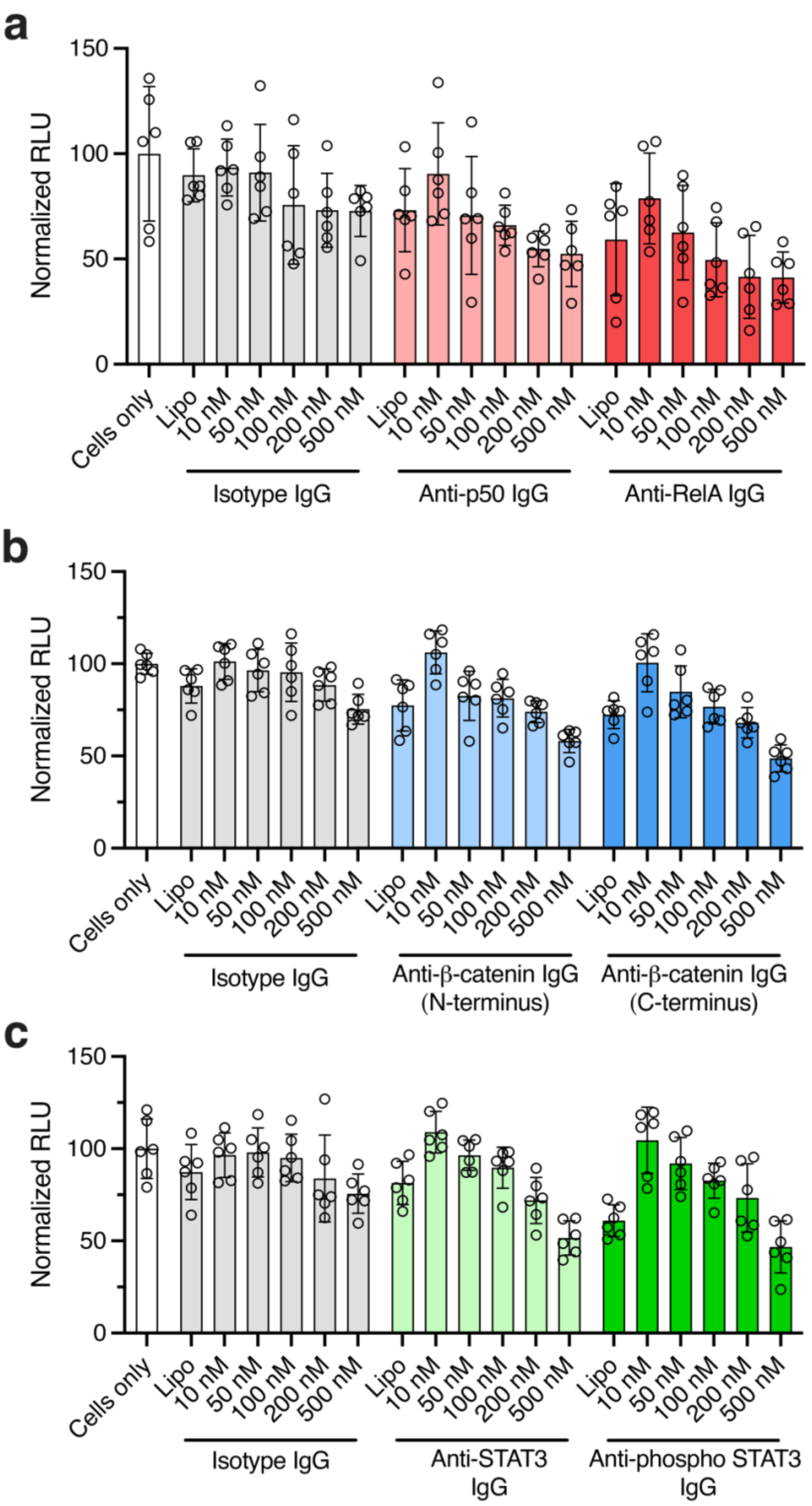


**Supplementary Figure S9. Signaling inhibition from luminescence assays of cells treated with IgG-encapsulated LNPs.** Data shown are for IgGs cloaked with 30 molar eq. of SL4 and for MC3 LNPs (MC3/IgG, 2 wt/wt) supplemented with 30 mol % DOTAP and formulated in pH 5 PBS. All transfections were performed for 12 hours. (a-c) Luminescence readouts of transcriptional activity following 200 nM transfections of cloaked IgGs with LF2K and 10 nM – 500 nM transfections of cloaked IgGs with MC3 LNPs of (a) anti-RelA and anti-p50 IgGs against NF-κB signaling in A549 cells, (b) N-terminal and C-terminal anti-β-catenin IgGs against Wnt signaling in DLD-1 cells, and (c) anti-STAT3 and anti-phospho STAT3 IgGs against JAK-STAT signaling in HepG2 cells, along with isotype control IgGs. All data are presented as mean ± SD (*n* = 6 for luminescense assays).

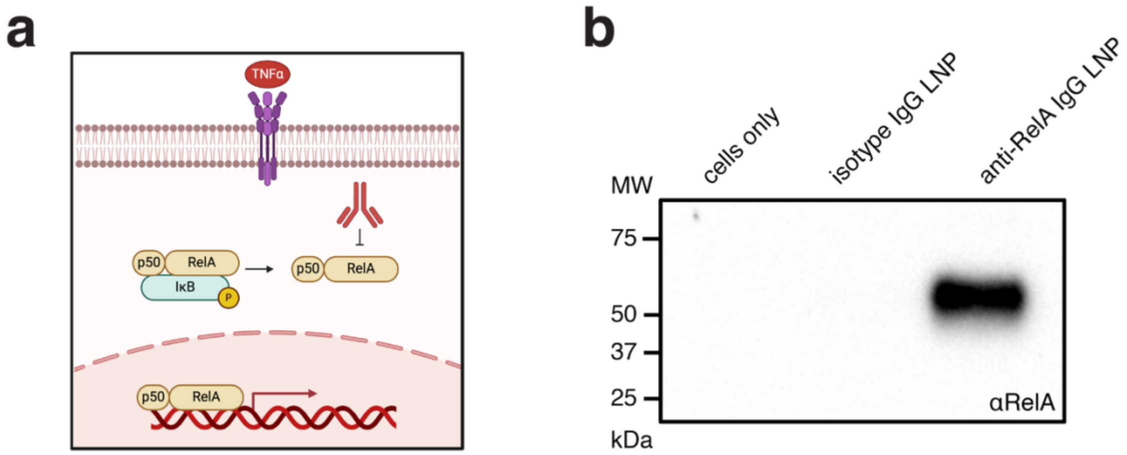


**Supplementary Figure S10. Co-immunoprecipitation (co-IP) of anti-RelA IgGs bound to RelA in A549 cells.** (a) Schematic depicting RelA inhibition with anti-RelA IgGs in NF-κB pathway. (b) Western blot against RelA following protein G pull-down of transfected anti-RelA IgGs from A549 cell lysates. IgGs were cloaked with 30 molar eq. of SL4 formulated with MC3 LNPs (MC3/IgG, 2 wt/wt) supplemented with 30 mol % DOTAP in pH 5 PBS.

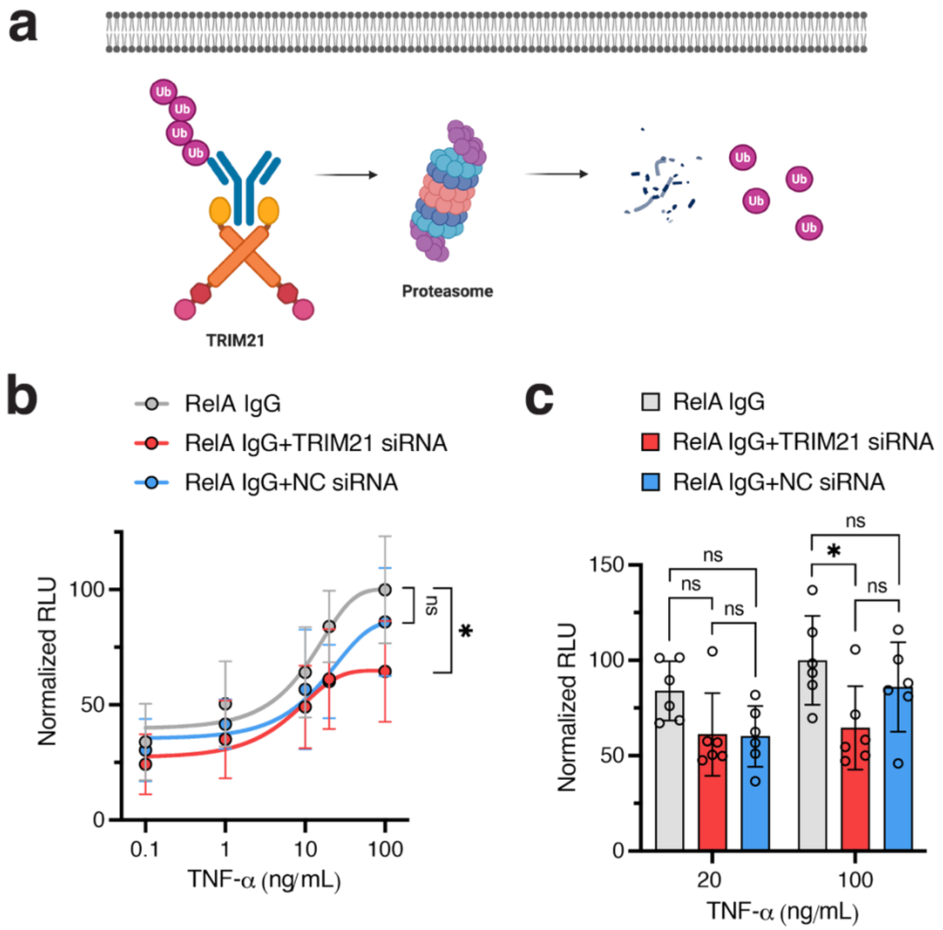


**Supplementary Figure S11. Effects of TRIM21 on inhibition of NF-κB inhibition following delivery of IgG-encapsulated LNPs.** Data shown are for IgGs cloaked with 30 molar eq. of SL4 and for MC3 LNPs (MC3/IgG, 2 wt/wt) supplemented with 30 mol % DOTAP and formulated in pH 5 PBS. All transfections were performed for 12 hours. (a) Schematic depicting TRIM21-mediated degradation of cytosolic antibodies. (b) Luminescence readouts of NF-κB transcriptional activity following co-transfections of 100 nM of cloaked anti-RelA IgGs and either TRIM21 siRNA or negative control (NC) siRNA with MC3 LNPs in A549 cells. Cells were stimulated with varying concentrations of TNF-α (0.1 – 100 ng/mL) for 5 hours before luminescence readings. (c) Comparison of NF-κB transcriptional activity following co-transfections in A549 cells at 20 ng/mL and 100 ng/mL of TNF-α stimulation. Data are presented as mean ± SD ($n$ = 6 for luminescence assays). Statistical significance determined by two-way ANOVA and one-way ANOVA followed by Bonferroni correction for multiple comparisons (*$p < 0.05$, **$p < 0.01$, ***$p < 0.001$, ****$p < 0.0001$).

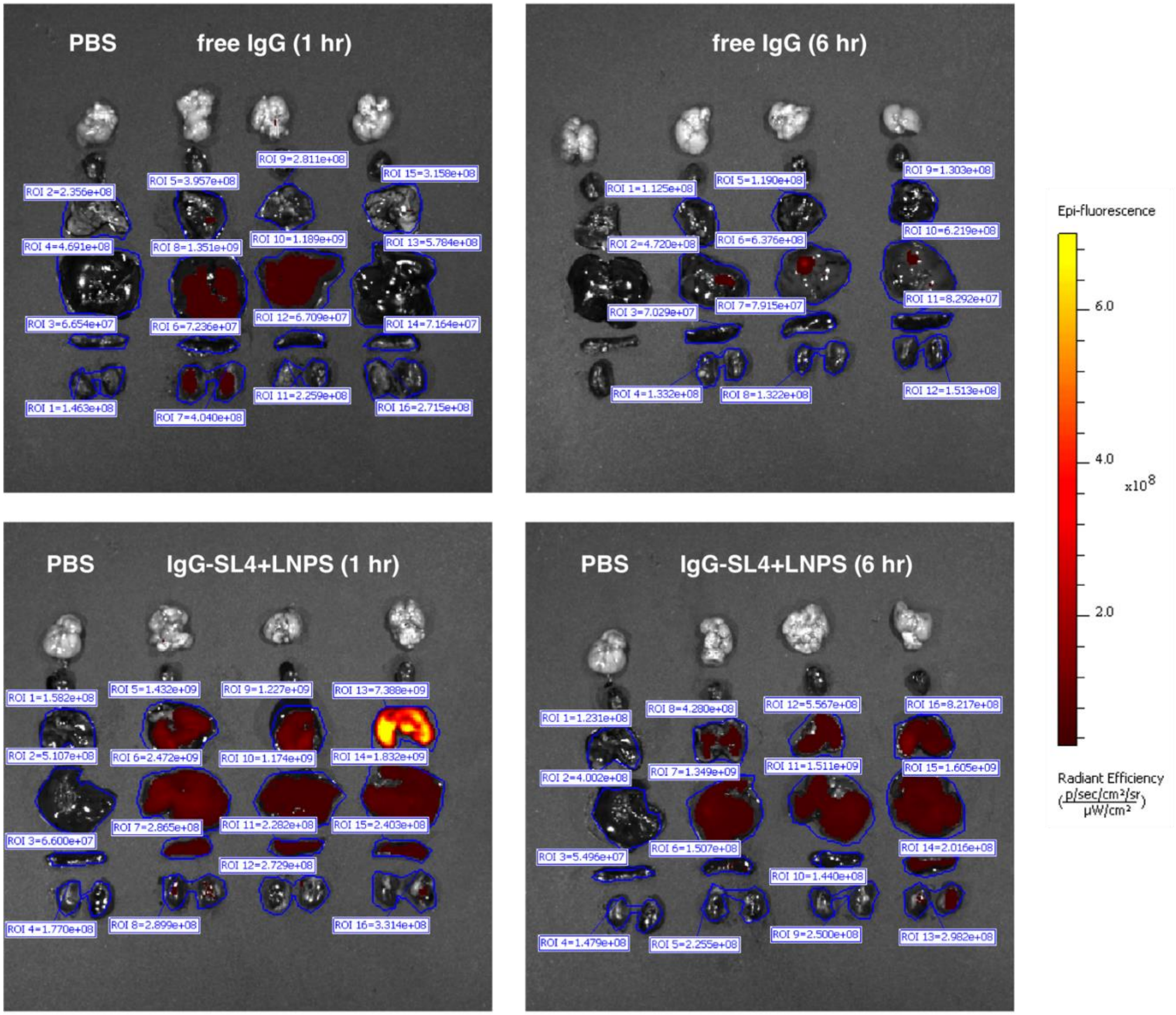


**Supplementary Figure S12. *In vivo* biodistribution of cloaked-IgGs delivered with LNPs.** *Ex vivo* fluorescent images of harvested organs following tail vein injections of BALB/c mice with PBS, free IgG, and IgG-SL4 formulated in MC3 LNPs. MC3 LNPs were formulated with 3 mol% PEG-DMG-2000 and 30 mol% DOTAP. Doses were 1 mg/kg of total protein. Images were taken 1 h and 18 h post-injection. Calculated radiant efficiencies of ROIs corresponding to lungs, liver, and kidneys are shown.

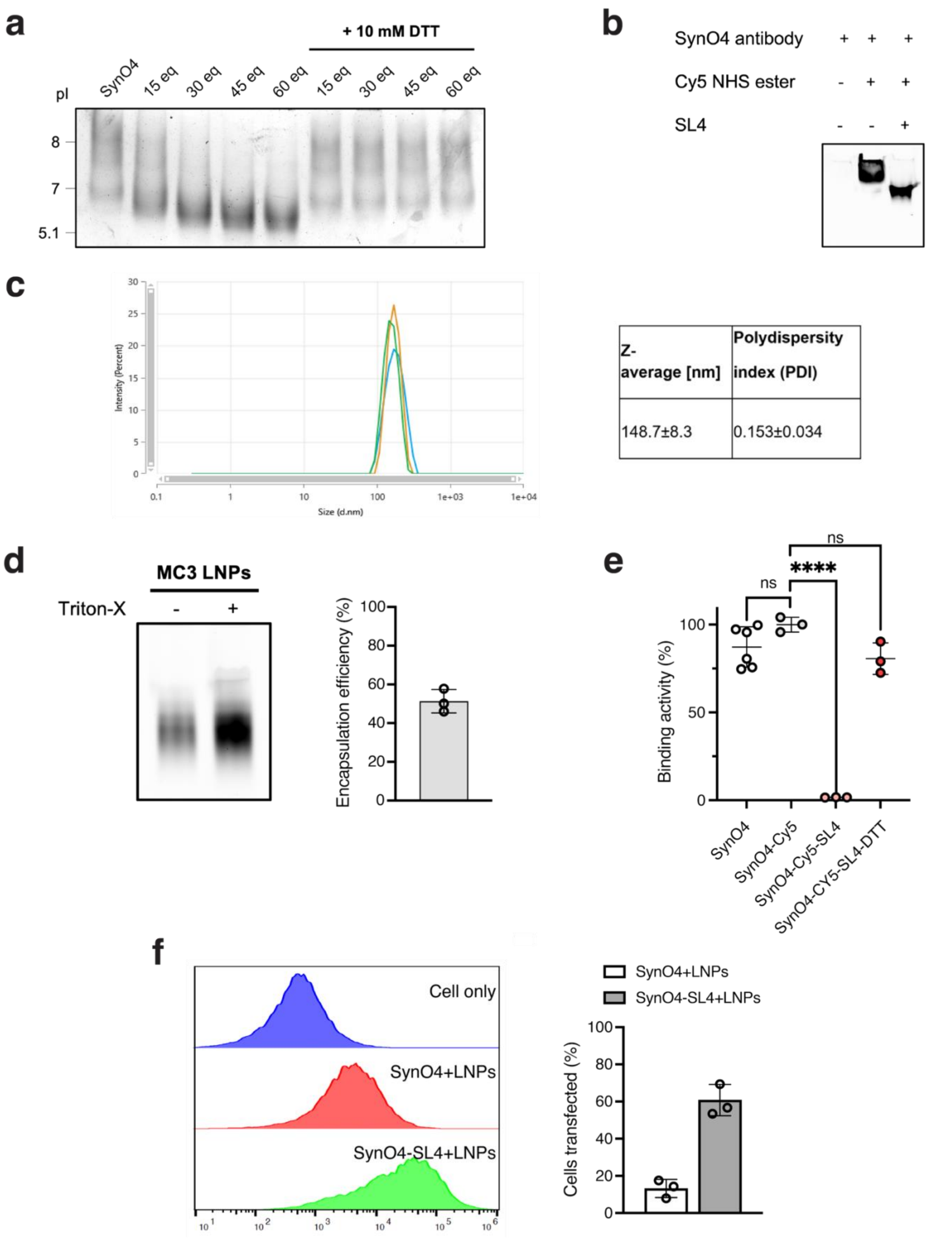


**Supplementary Figure S13. Formulation and characterization of SynO4-loaded LNPs.** (A) Isoelectric focusing (IEF) gels of SynO4 monoclonal antibody conjugated to SL4, before and after treatment with 10 mM DTT (b) In-gel fluorescence of SynO4, SynO4-SL4 and SynO4-SL4-Cy5 under native conditions (c) Representative dynamic light scattering (DLS) profile of SynO4-Cy5-LNPs. Average particle size (Z-average) and polydispersity index (PDI) are shown in the accompanying table. (d) In-gel fluorescence of SynO4-Cy5-LNPs under native conditions, with or without Triton X-100 treatment; the right panel quantifies encapsulation efficiency of SynO4-SL4-Cy5 in MC3 LNPs based on densitometry analysis. (e) Binding activity of SynO4 samples to immobilized alpha-synuclein as determined by ELISA, in the presence or absence of DTT; data are presented as percent activity relative to native form of SynO4 (f) Representative flow cytometry histograms of HEK293T cells transfected with 200 nM of SynO4 IgGs and SynO4 IgGs cloaked with SL4 using MC3 LNPs. All transfections performed for 6 hours; the right panel quantifies cell uptake. All data are presented as mean ± SD (*n* = 3 native gels; n = 3 – 6 for ELISA; n = *3* for flow cytometry). Statistical significance was determined by one-way ANOVA followed by Bonferroni correction for multiple (*p < 0.05, **p < 0.01, ***p < 0.001, ****p < 0.0001).

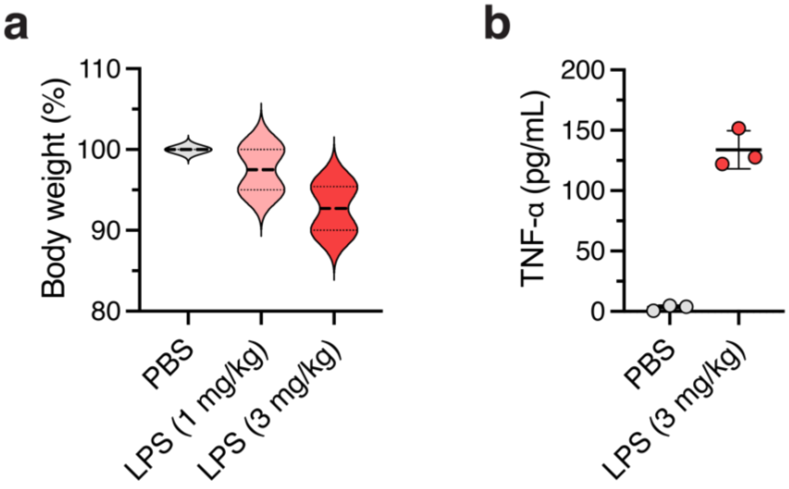


**Supplementary Figure S14. Establishment of acute lung injury model in BALB/c mice.** (a) Body weight change of BALB/c mice 24 hours after intratracheal instillation with PBS and LPS. (b) Cytokine levels of TNF-α and from extracted BALF samples in mice 24 hours after PBS and LPS instillation. All data are presented as mean ± SD (*n* = 3 for mice injections).

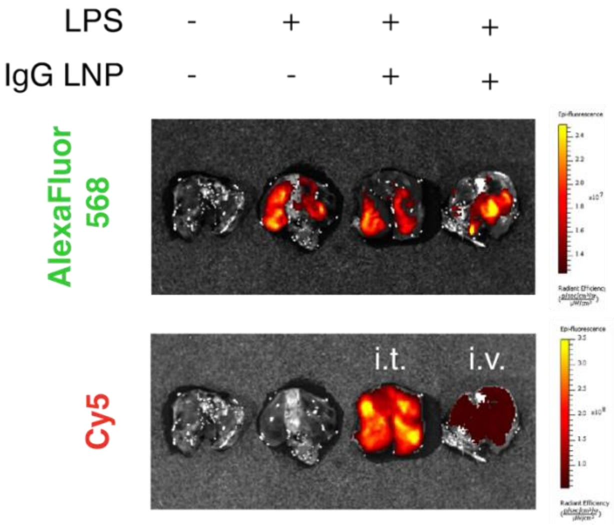


**Supplementary Figure S15. Localization of fluorescently labeled LPS and cloaked IgGs in LPS *in vivo*.** *Ex vivo* fluorescent images of harvested lungs following intratracheal injections of AlexaFluor 546-labeled LPS along with intratracheal (i.t.) and intravenous (i.v.) injections of Cy5-labeled cloaked IgGs in MC3 LNPs into BALB/c mice. Images shown are for 1 hour post-injection.

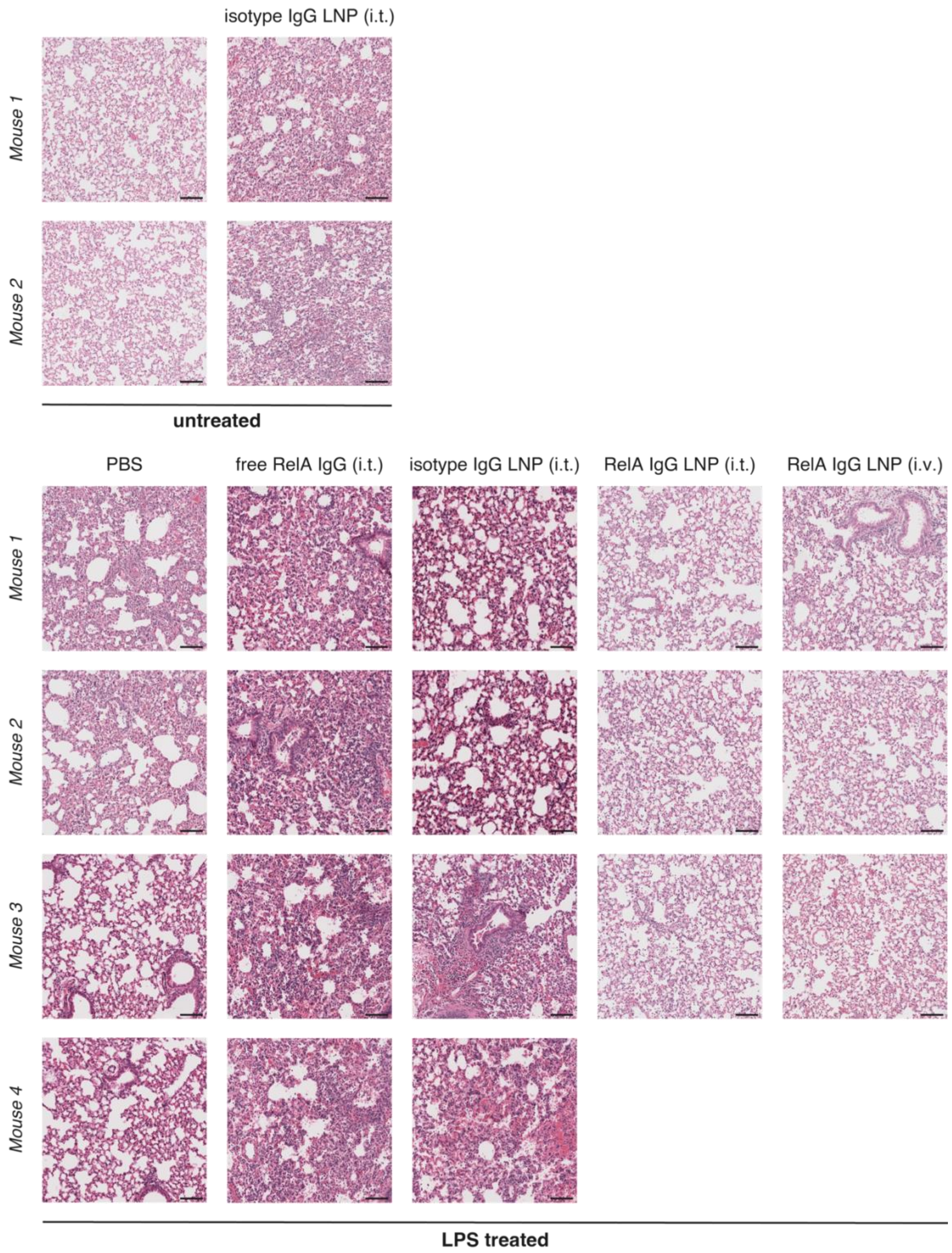


**Supplementary Figure S16. Histological analysis of sectioned lung tissue from ALI-induced mice.** Representative images of H&E-stained lung tissue samples obtained from BALB/c mice following IgG treatments. Scale bar = 100 μm.

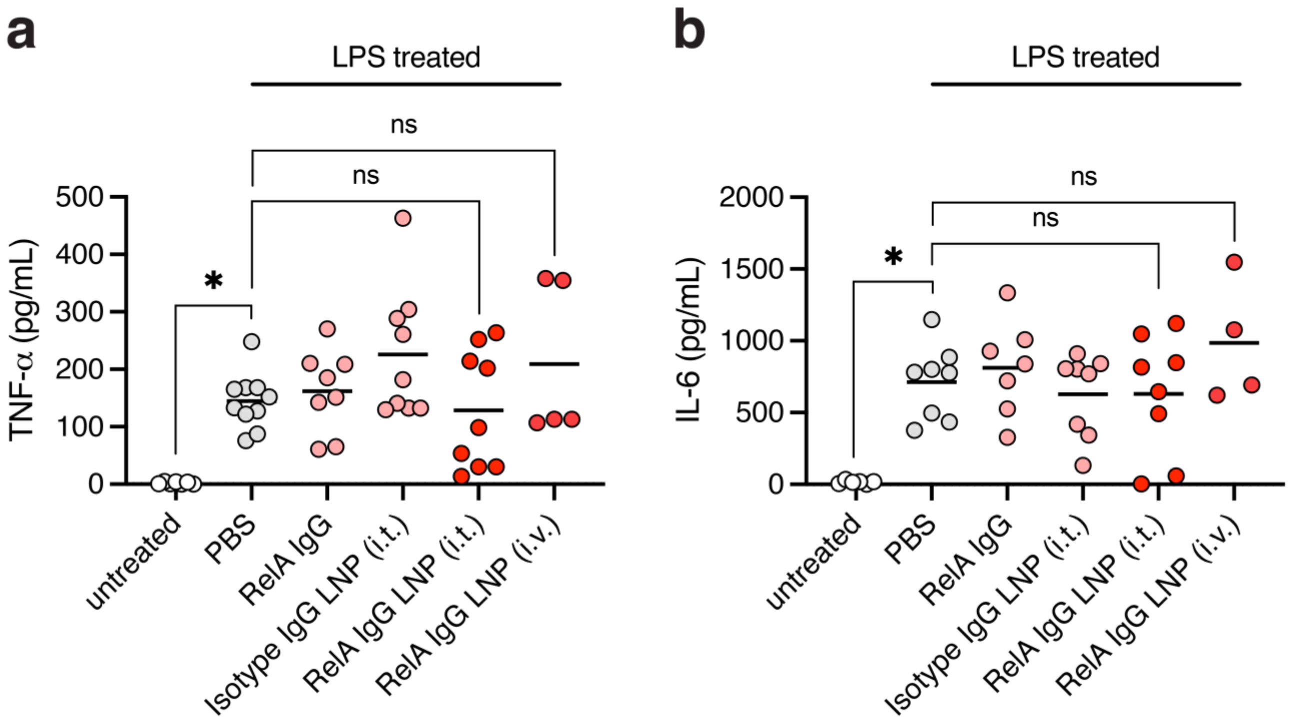


**Supplementary Figure S17. ELISA of pro-inflammatory cytokines from harvested bronchioalveolar lavage fluid (BALF) from ALI-induced mice.** Cytokine levels of (a) TNF-α and (b) IL-6 from extracted BALF samples in mice following IgG treatments. All data are presented as mean ± SD (*n* = 4 – 8 for ELISA). Statistical significance was determined by one-way ANOVA followed by Bonferroni correction for multiple (*p < 0.05, **p < 0.01, ***p < 0.001, ****p < 0.0001).

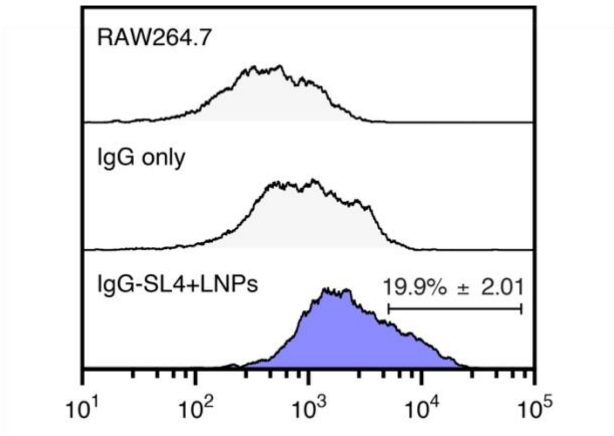


**Supplementary Figure S18. Flow cytometry results of RAW264.7 cells treated with IgG-encapsulated LNPs using fluorogen activating protein assay.** Representative flow cytometry histograms of RAW264.7 cells transfected with 200 nM of IgGs and IgGs cloaked with SL4 using MC3 LNPs. Data shown are for IgGs cloaked with 30 molar eq. of SL4 and for MC3 LNPs (MC3/IgG, 2 wt/wt) supplemented 30 mol % DOTAP and formulated in pH 5 PBS. All transfections performed for 6 hours.

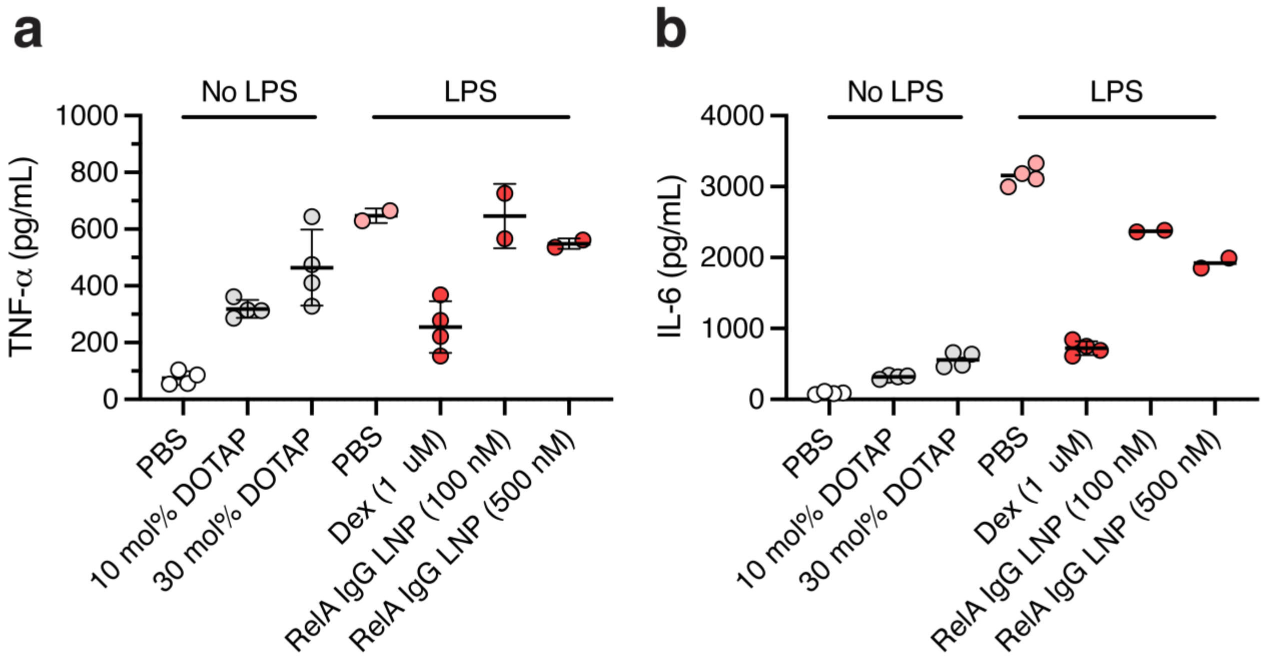


**Supplementary Figure S19. ELISA of pro-inflammatory cytokines from RAW264.7 cells.** Cytokine levels of (a) TNF-α and (b) IL-6 from unstimulated and LPS-stimulated RAW264.7 cells following IgG and dexamethasone (Dex) treatments. Data shown are for IgGs cloaked with 30 molar eq. of SL4 and for MC3 LNPs (MC3/IgG, 2 wt/wt) supplemented 30 mol % DOTAP and formulated in pH 5 PBS. All data are presented as mean ± SD (*n* = 2 – 4 for ELISA).

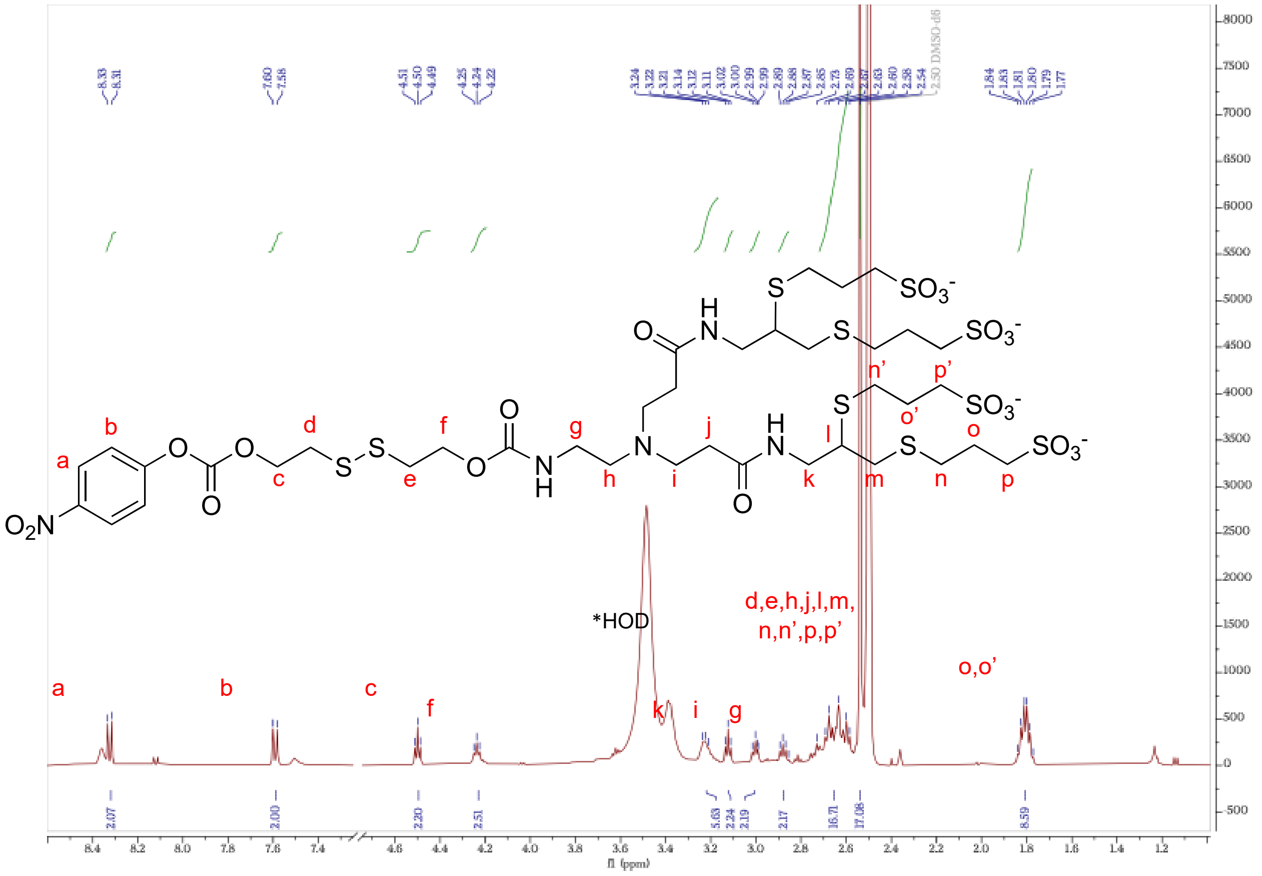


**Supplementary Figure S20.** $^1$H NMR (500 MHz, DMSO-d$_6$) of SL4.

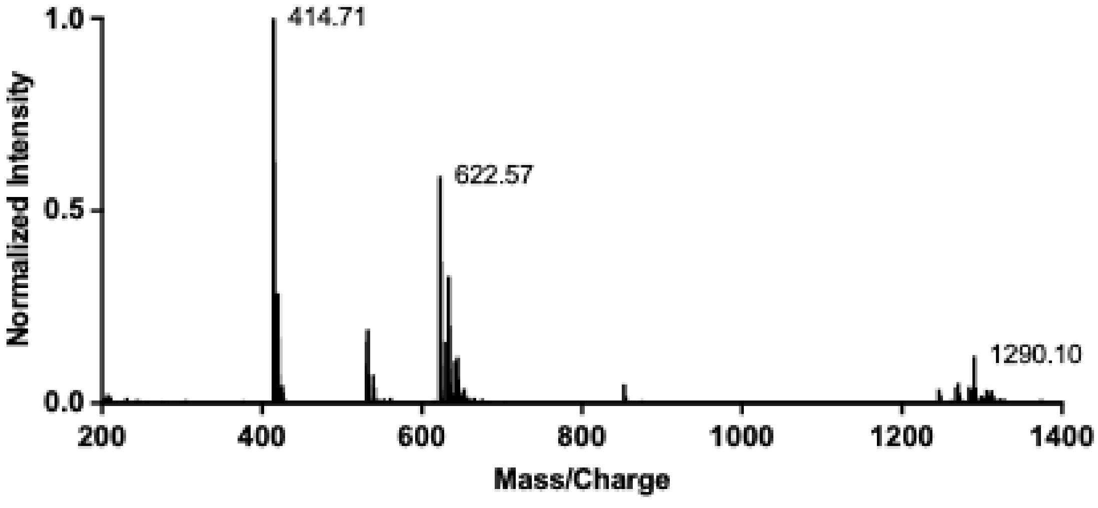


**Supplementary Figure S21.** LC/MS analysis of SL4 (negative ion mode).
